# Restoration of DNA Integrity and Cell Cycle by Electric Stimulation in Planarian Tissues Damaged by Ionizing Radiation

**DOI:** 10.1101/2021.05.16.444365

**Authors:** Devon Davidian, Melanie LeGro, Paul G. Barghouth, Salvador Rojas, Benjamin Ziman, Eli Isael Maciel, David Ardell, Ariel L. Escobar, Néstor J. Oviedo

**Affiliations:** Department of Molecular & Cell Biology, University of California, Merced, USA.; Quantitative and Systems Biology Graduate Program, University of California, Merced, USA.; Department of Bioengineering, University of California, Merced, USA.; Health Sciences Research Institute, University of California, Merced, USA.

**Author notes:** To whom correspondence should be addressed: Néstor J. Oviedo, Department of Molecular & Cell Biology, University of California. 5200 North Lake Road, Merced, CA 95343. Equal contribution (Devon Davidian, Melanie LeGro).

**Keywords:** electric stimulation, stem cells, DNA repair, planaria, neoblasts, tissue regeneration, galvanotactic

## Abstract

Exposure to high levels of ionizing γ-radiation leads to irreversible DNA damage and cell death. Here, we establish that exogenous application of electric stimulation enables cellular plasticity to reestablish stem cell activity in tissues damaged by ionizing radiation. We show that sub-threshold direct current stimulation (DCS) rapidly restores pluripotent stem cell populations previously eliminated by lethally γ-irradiated tissues of the planarian flatworm *Schmidtea mediterranea*. Our findings reveal that DCS enhances DNA repair, transcriptional activity, and cell cycle entry in post-mitotic cells. These responses involve rapid increases in cytosolic [Ca^2+^] through the activation of L-type Cav channels and intracellular Ca^2+^ stores leading to the activation of immediate early genes and ectopic expression of stem cell markers in postmitotic cells. Overall, we show the potential of electric current stimulation to reverse damaging effects of high dose γ-radiation in adult tissues. Furthermore, our results provide mechanistic insights describing how electric stimulation effectively translates into molecular responses capable of regulating fundamental cellular functions without the need for genetic or pharmacological intervention.

## INTRODUCTION

Since the initial observations made by Luigi Galvani in the late 1700s, scientists have been fascinated by the effects of the exogenous application of electric currents in animal tissues (Adee, 2018; Bresadola, 1998). This long-lasting interest has revealed almost universal roles for electricity during embryonic development, tissue regeneration, and disease (Levin, 2007; Levin, 2014; McCaig et al., 2005). For example, modern FDA approved applications of electric stimulation include deep brain stimulation where indwelling electrodes are implanted in specific regions of the brain to deliver electrical currents and treat diseases such as essential tremors, slow movement, and stiffness in Parkinson disease (Graupe et al., 2018; Miocinovic et al., 2013; Mohammed et al., 2018; Niemann et al., 2017; Velarde et al., 2017). Electric stimulation is also used in treatment-resistant (i.e., refractory) depression, which is the leading cause of disability worldwide (Bewernick et al., 2012; Delaloye and Holtzheimer, 2014; Holtzheimer et al., 2012; Mayberg et al., 2005; Organization, 2018). Recent studies with electric stimulation demonstrate the possibility to restore voluntary control of walking in animals and humans with spinal cord injury (Borgens et al., 1987; Formento et al., 2018; Wagner et al., 2018). Extensive clinical trials, over the past 50 years, demonstrate that direct current stimulation (DCS) improves bone healing by accelerating repair time and reducing pain in nonunion bone fractures (Aleem et al., 2016; Brighton, 1981; Brighton et al., 1981; Griffin and Bayat, 2011; Kuzyk and Schemitsch, 2009; Zhuang et al., 1997).

Additionally, DCS directly applied to the surface of the scalp, (i.e., without implanting electrodes) is known as transcranial DCS (tDCS) and is commonly used in humans (Antal et al., 2017; Chaieb et al., 2014; Chang et al., 2016; Elliott, 2014; Huang et al., 2015b; Kadosh, 2015; Kadosh et al., 2010; Moreno-Duarte et al., 2014; Nelson et al., 2016; Sarkar and Kadosh, 2016; Tortella et al., 2015). tDCS is widely used to treat many human conditions including motor learning in stroke neurorehabilitation (Boggio et al., 2007), migraines (Antal et al., 2011; Antal et al., 2008), patients with chronic pain (Boggio et al., 2009; Boggio et al., 2008; Borckardt et al., 2012; Mylius et al., 2012), and psychiatric disorders (Tortella et al., 2015). Likewise, cognitive assessments have shown that tDCS can be used to modulate the rate of learning and improve numerical competence in humans (Kadosh, 2015; Kadosh et al., 2010; Snowball et al., 2013). Despite the widespread use of DCS in experimental, clinical, and private settings, the molecular bases of its effects in various cell types remain largely unknown. Together with the rapid proliferation and easy access to devices marketed as direct to customer technology involving electrical stimulation has raised ethical questions, produced confounding results, and created concerns about the risk of potential adverse effects in the body (Coates McCall et al., 2019; Wexler and Reiner, 2019; Wurzman et al., 2016).

The effects of DCS vary depending on the location in the body, the tissue type that is targeted, the intensity of the electric current, and the duration of treatment. Furthermore, the outcome of DCS may differ depending on the context in which it is assayed (i.e., embryonic development, adult stages, disease or homeostasis). It is important to note that currents used in DCS are subthreshold, implying they do not induce action potentials in excitable cells/tissues (Bikson et al., 2004). All cells, including stem cells (SCs), progenitors, and differentiated cells, have built-in mechanisms designed to sense electric changes, which consequently influence their innate cellular transmembrane potential (V_mem_). Changes in V_mem_ are caused by changes in ion fluxes at the cell plasma membrane (Jaffe, 1981a; Jaffe, 1981b; Levin, 2007; McLaughlin and Levin, 2018; Nuccitelli and Jaffe, 1974). V_mem_ acts as a potent regulator of cellular migration, proliferation, differentiation, cell cycle, and cell death and is highly sensitive to manipulation by DCS (Levin, 2014; McCaig et al., 2005). Furthermore, DCS, at the organismal level, is capable of altering axial polarity, tissue repair, and organ specification (Borgens et al., 1987; Jaffe, 1981a; Marsh and Beams, 1952; McCaig et al., 2005; McLaughlin and Levin, 2018; Nuccitelli, 2003; Zhao et al., 2006). For example, galvanotactic responses have been observed in migratory cells of humans (Guo et al., 2010), fish (Graham et al., 2013), frogs (Stump and Robinson, 1983), *C. elegans* (Chrisman et al., 2016), and even Paramecium (Ogawa et al., 2006). Similarities in DCS-mediated cellular responses across vertebrate and invertebrate organisms argue for evolutionarily conserved mechanisms.

To further investigate the effects of DCS, we developed an experimental strategy using the planarian flatworm *Schmidtea mediterranea*, known for their high rates of cellular turnover and extraordinary capacity to regenerate tissues that rely in adult SCs called neoblasts (Reddien, 2018; Rink, 2018; Zeng et al., 2018b; Zhu and Pearson, 2016). Neoblasts are known as the only cell with the capacity to divide in asexually reproducing planarians. Thus, neoblast division alone provides the cellular progeny required to renew and repair all planarian tissues (van Wolfswinkel et al., 2014b; Wagner et al., 2011; Zeng et al., 2018b). This constant cellular crosstalk allows planaria to maintain a diverse cellular population through neoblast division, making them powerful models to analyze the effects of DCS on adult SCs and differentiated cells at the cellular, subcellular, and organismal level. Here, we introduced a simplified platform to apply exogenous DCS to the whole body of planarians and analyze the resulting cellular and subcellular responses in real-time. We found that brief exposure to DCS overrides cellular decisions in tissues exposed to lethal doses of ionizing radiation. Furthermore, our results reveal that DCS is a rapid and robust method with the potential to enhance DNA damage repair, activate transcription of stem cell markers in tissues damaged by ionizing radiation. Moreover, we identified that these DCS-mediated responses are tightly regulated by transcription of immediate early genes and rapid intracellular Ca^2+^ flux. These findings provide insights into the effects of DCS in the adult body, exhibiting their domain over fundamental cellular processes such as transcription, DNA repair, cell cycle, and cellular plasticity without the need for genetic or pharmacological treatments.

## RESULTS

### Body-wide Application of Steady-State Direct Current (pDCS)

We immobilized planarians by applying our recently developed method (i.e., ThermoPress immobilization), which combines the anesthetic chloretone (0.2%) with a 1% agar-encasing chamber (Davidian et al., 2020). This method of agar immobilization keeps planarians alive while preserving tissue integrity and restricting body movement for hours to days with complete and rapid recovery of sensory functions and locomotion (Fig. 1A). ThermoPress immobilization was used to administer steady-state direct current stimulation to the whole planarian body, which we coined ‘pDCS’ (Davidian et al., 2020).

**Fig. 1.**
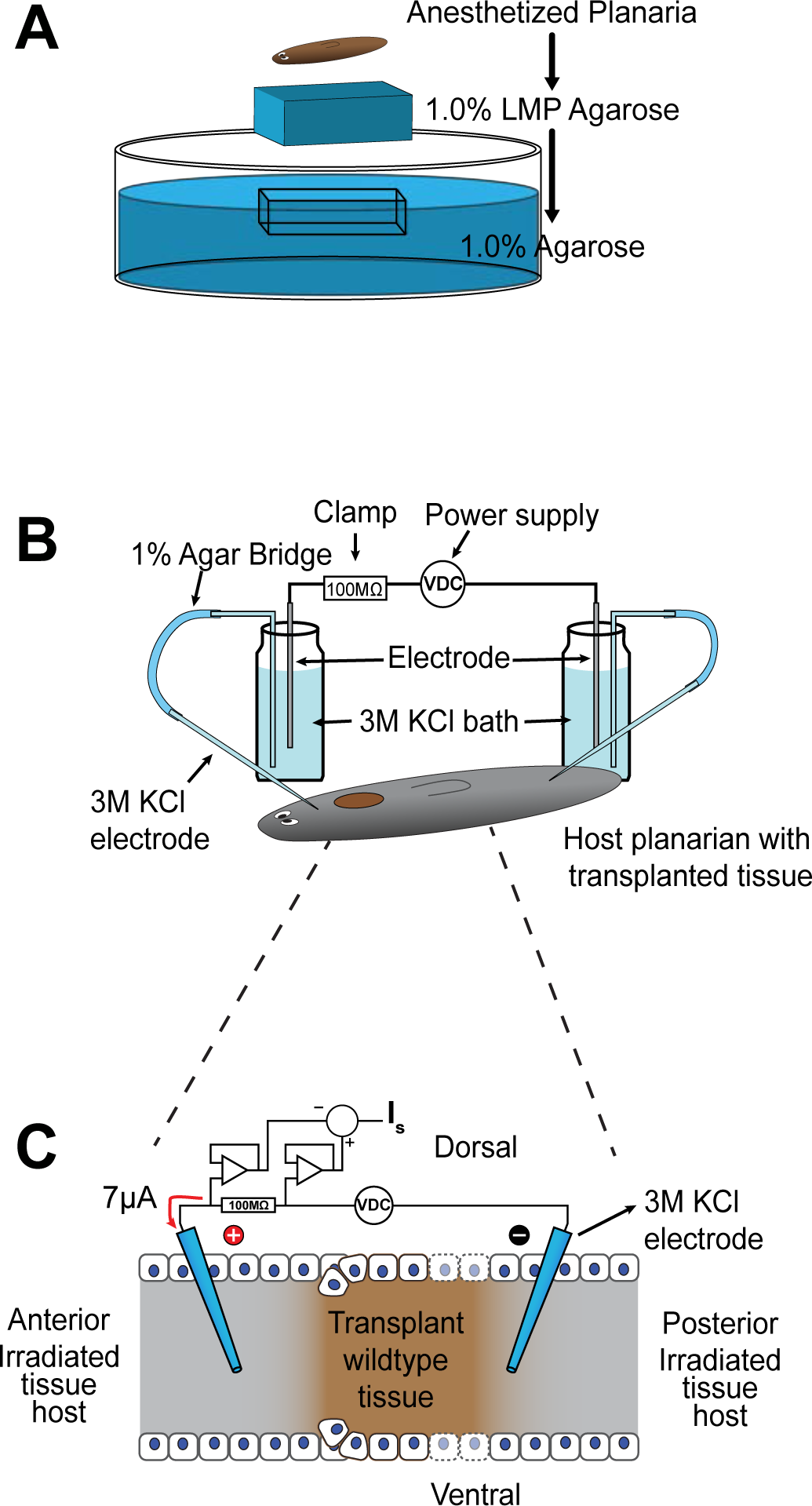
Schematic summary of planarian immobilization and pDCS setup. (A), Schematic representation of planarian immobilization. (B), Illustration of current-clamp circuit created by pDCS electrodes and planarian. (C), Amplified schematic representation of planarian tissues showing electrode placement and op-amps used to quantify current passing through the 100MΩ current clamp.

Electric current was delivered through pulled borosilicate sharp microelectrodes, filled with 3M KCl, placed at the anterior and posterior ventral side of planarian (i.e., pre-pharyngeal and tail, respectively). The microelectrodes resistance was consistently measured at 1-2MΩ. The control group (sham-treated), consisted of animals exposed to the same procedure, included electrode penetration, with the absence of electric current. To facilitate circuit conduction and minimize electrode byproduct in planarian tissues, microelectrodes were coupled to an electrode bath using 1.0% agar bridges containing 3M KCl solution (Fig. 1B). The amount of current delivered was limited through a 100MΩ resistor, and the most consistent results were obtained when the current traversing the animal was 7µA (Fig. 1C). DCS exceeding 7µA resulted in noticeable tissue damage and eventual animal lysis. All experiments were performed with an electric polarity of a positive pole in the anterior and negative pole in the posterior, unless otherwise stated. Current delivered to the animal was differentially measured and acquired with an analog-to-digital converting board controlled with custom-made LabView-based software (Elliott et al., 2007). Animals were under constant surveillance to ensure that electrode placement and tissue integrity were maintained. Overall, this setup was effective in keeping electrodes correctly positioned and allowed for the analysis of DCS for up to 6 hours.

### pDCS activates transcription of stem cell markers in tissues exposed to a lethal dose of ionizing radiation

DCS techniques are known to affect both SCs and differentiated cells in many model organisms (Feng et al., 2017; Huang et al., 2015a; McCaig et al., 2005; Zhao et al., 2006). Planarian neoblasts are generally scattered along the antero-posterior (AP) axis except for regions in front of the eyes or pharynx and uniquely express the gene *smedwi-1* (Reddien et al., 2005; van Wolfswinkel et al., 2014b; Wagner et al., 2011; Zeng et al., 2018b) (*piwi-1* henceforth, Fig. 2A). The *piwi-1* expression is currently used as the standard marker to recognize neoblast presence and distribution (Reddien et al., 2005; Wagner et al., 2011; Zeng et al., 2018b).

**Fig. 2.**
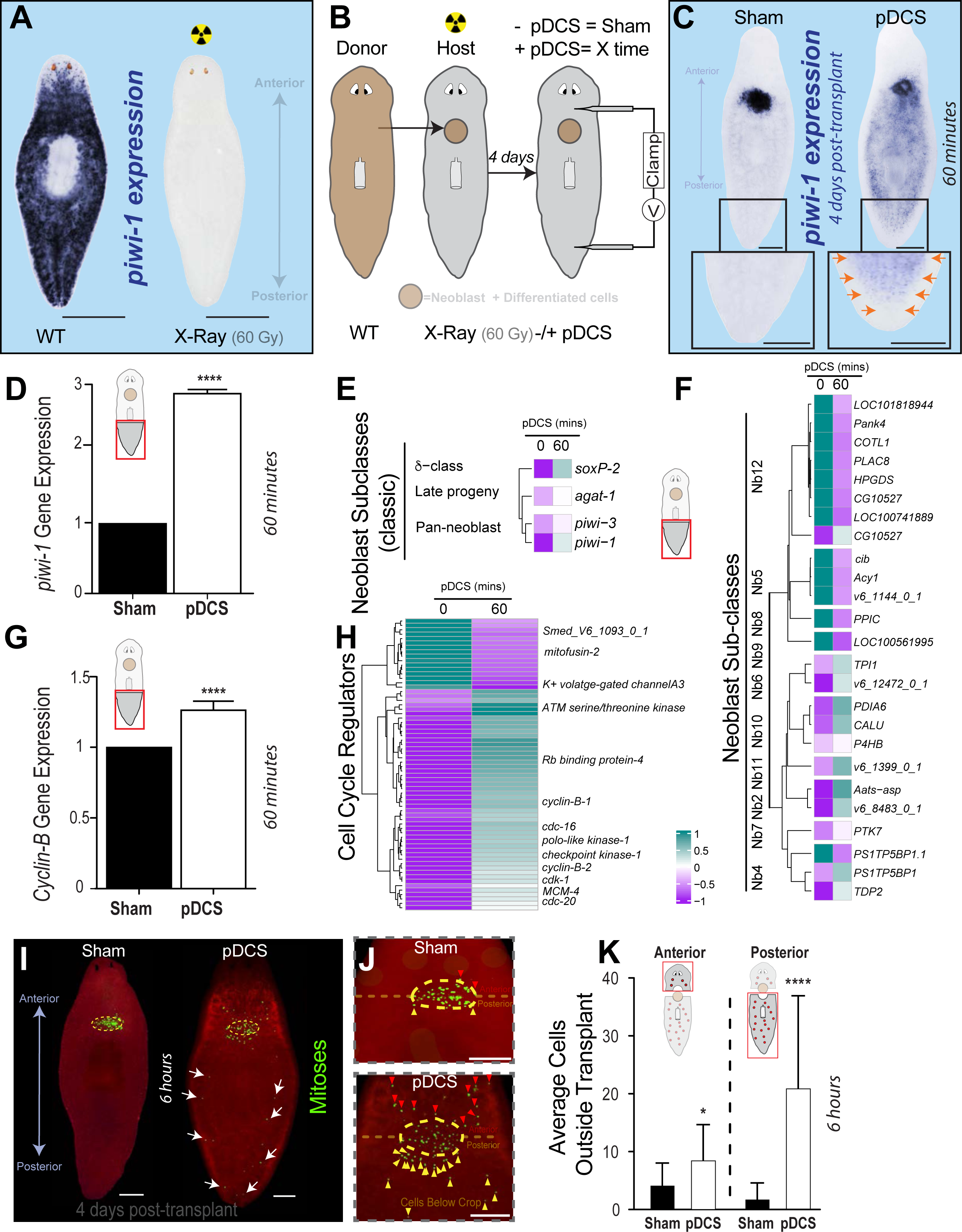
pDCS triggers a transcriptional response and cell cycle in γ-irradiated host tissues. (A) Whole-mount *in situ* hybridization (WISH) showing *piwi-1^+^* signal in both wild-type (WT) and γ-irradiated (γ-irr, 60Gy) planarian (n=10/10). (B) Schematic depicting transplantation procedure using WT donor and irradiated host planarian with subsequent exposure to DCS. (C) WISH of *piwi-1* gene expression after four days post-transplant in both sham (control, n=10/10) and animals subjected to 60min pDCS (n=12/15). The insets in the lower portion of the image represent magnification of the distal part of the animal –tail-, *piwi-1* signal is indicated with arrows in animals subjected to pDCS. *(D) piwi-1* gene expression levels as determined by qPCR (pool of six animals/replicate and three biological replicates). (E-F) RNA-seq data collected from the host-tail tissue of sham and animals subjected to 60 mins pDCS (data was collected by pooling tails from four independently treated pDCS or sham planaria across three independent experimental trials). Gene expression heatmaps display differentially expressed transcripts (FDR<0.05) as averaged log_2_CPM Z-scores. (E) Heatmap representation of RNA-seq data displays differentially expressed neoblast subclass populations using the classical neoblast classification from van Wolfswinkel et al., 2014b; Wagner et al., 2012. (F) RNA-seq heatmap displays the neoblast subpopulations and their respective lineages based on recent neoblast classification (Zeng et al., 2018b). (G) *cyclin-B* gene expression levels obtained with qPCR from tail fragments in both the sham and 60min pDCS (pool of six animals/replicate and three biological replicates). (H) RNA-seq heatmap displaying differentially expressed genes commonly associated with cell cycle regulation. (I) Whole-mount immunostaining with anti-H3P antibody (green dots) showing H3P^+^ cells inside and outside of the transplant. Notice H3P^+^ cells in the experimental group far away from the transplanted tissue (white arrows) following 6hrs pDCS compared to sham control (n=10/15). Dotted yellow circle: transplanted tissue (J) Magnified images around the transplanted tissue for both sham and pDCS after 6 hours of treatment (red and yellow arrows indicate mitotic cells in the anterior and posterior to the transplant -pink dotted line). (K) Average of H3P^+^ cells in sham and the experimental groups after 6 hours of pDCS. pDCS experiments were executed with positive pole to the anterior and negative to the posterior for 60 minutes. Data represented as mean ± SEM. Students *t*-test: \*\*\*\**P* ≤ 0.0001. Scale bars, 500µm.

Exposing planarians to lethal doses of ionizing radiation (60 Gy) irreversibly eliminates neoblasts and corresponding *piwi-1* expression (Fig. 2A); consequently, abolishing their regenerative capabilities. As a result, planarians ultimately perish within three to four weeks following lethal ionizing radiation (Bardeen and Baetjer, 1904; Reddien et al., 2005). However, lethally irradiated planarians can be rescued by transplanting tissue-containing neoblasts (Guedelhoefer and Sanchez Alvarado, 2012a; Guedelhoefer and Sanchez Alvarado, 2012b). This occurs as neoblasts gradually migrate from transplanted tissues to repopulate the entire irradiated host in about a month (Guedelhoefer and Sanchez Alvarado, 2012a). Engrafted tissue becomes more structurally stable after four days post-transplantation (dpt); a stage in which the majority of neoblasts are still within the transplanted tissue. Over the first five dpt, neoblast repopulation is uniform with no bias towards anterior, posterior, medial-lateral tissues (Guedelhoefer and Sanchez Alvarado, 2012a). Thus, this neoblast repopulation paradigm was used as a model to study the effects of pDCS on both adult SCs and differentiated cells in their natural environment.

Tissue containing neoblasts was transplanted into lethally irradiated hosts and pDCS-treated at four dpt (Fig. 2B). This experimental setup led to essential differences in *piwi-1* gene expression in sham vs. pDCS animals. Specifically, 60 minutes after pDCS, *piwi-1* expression was widely detected outside the transplant toward the posterior region of the animal; whereas, in the sham-treated group, *piwi-1* expression was restricted to tissues adjacent to the transplant, as expected (Fig. 2C). Levels of *piwi-1* expression in the tails of treated planarian increased by more than two-fold as determined by using qPCR (Fig. 2D). Transcriptomic analysis with RNA sequencing from the tail fragment confirmed pDCS treatment for 60 mins was accompanied by an increase in the expression of *piwi-1* (Log2 FC= 1.97, B&H FDR= 0.0006, moderated t= 9.89).

Furthermore, we also detected an increase in the expression of the transcription factor *Smed-soxp-2* (Fig. 2E, Log2 FC= 0.61, B&H FDR= 0.005, moderated t= 4.82), which is a marker of the classic sigma neoblast subpopulation required for stem cell function and planarian regeneration (van Wolfswinkel et al., 2014a; Wagner et al., 2012). Likewise, we observed pDCS slightly reduced the expression of the *piwi* family member *Smed-piwi-*3 after 60 mins of pDCS (Log2 FC= 0.21, B&H FDR= 0.0068, moderated t= 4.24) (Kim et al., 2019; Palakodeti et al., 2008b), while there was moderately upregulation of the late progeny marker *Smed-agat-1* (Fig. 2E, Log2 FC= 0.18, B&H FDR= 0.016, moderated t= 3.74).

We also observed differential expression of other markers (Fig. 2F) associated with the recently expanded neoblast sub-classes (Zeng et al., 2018a). Out of 189 possible Nb markers from the literature 25 markers were differentially expressed with a B&H FDR < 0.05. The transcriptomic analysis tested the contrasts between the 60-minute and sham controls evidencing 1,778 genes differentially expressed and all 25 of these markers are expressed below the 1% level. The transcriptomic analysis for the tissues treated with 60 minutes of pDCS showed key markers for the neoblast sub-classes Nb5, Nb8, and Nb12 were strongly downregulated compared to the control at this timepoint (Fig. 2F). Nb5 and Nb12 contain populations of *Piwi-1^high^* expressing cells and are hypothesized to include early progenitor cells for intestinal tissue (B&H FDR <0.05). Conversely, markers for the muscle cell progenitors Nb4 and Nb6 (e.g., TDP2, TPI1, and ACTB) were significantly upregulated (Fig. 2F, B&H FDR <0.01). Muscle progenitors are critical for providing positional information and are key drivers of tissue patterning during regeneration (Cote et al., 2019; Scimone et al., 2020; Scimone et al., 2017). Additionally, putative markers for the pharyngeal neoblast progenitors and differentiated populations (Nb7, Nb8, and Nb10) were overall differentially expressed. However, the markers *PTK7* and *PPIC* for the *Piwi-1^high^* expressing pharyngeal progenitor populations (Nb7 and Nb8) had lower expression compared to the *Piwi-1^low^* differentiated pharyngeal populations (Nb10) (B&H FDR <0.05). The markers *PDIA6*, *CALU*, and *P4HB* for the Nb10 neoblasts were significantly upregulated after hour-long application of pDCS (B&H FDR <0.05). Only one marker (dd_Smed_v6_1399_0_1) for the putative neural progenitor (Nb11) was significantly upregulated after 60 minutes of pDCS (Fig. 2F, Log2 FC= 0.34, B&H FDR= 40.043, moderated t= 3.71). Importantly, key markers for clonogenic neoblasts (cNeoblasts, Nb2) were significantly upregulated (e.g., *Aats-asp* and *soxP-2*; Fig. 2E, F, Log2 FC> 0.5, B&H FDR <0.01). Our findings show that the hour-long application of pDCS triggers the expression of the pan-neoblast marker *piwi-1* along with markers of neoblast subpopulations with high *piwi-1* expression in host tissues where neoblasts were permanently eliminated by exposure to a lethal dose of ionizing radiation.

### Lengthening pDCS leads to a directional cell cycle entry in tissues exposed to lethal ionizing radiation

pDCS-induced transcription of *piwi-1* in irradiated tissue was also accompanied by an increase in the expression of the neoblast marker *Smed-cyclin B* and other components associated with the regulation of cell division (e.g., cyclin-dependent kinases, mini-chromosome maintenance proteins-MCM-, checkpoint kinase, polo-like kinase, and Rb binding protein), (Fig. 2G, H). However, the presence of mitotic cells far away from the transplanted tissue was evident after lengthening pDCS to 6 hours (Fig. 2I, J). Strikingly, the number of mitotic cells outside of the transplant was significantly increased upon pDCS (Fig. 2K). In these animals, mitotic cells within irradiated tissues were primarily distributed towards the posterior region of the host, being observed as far as the tip of the tail (Fig. 2K). Proportionately, a smaller number of mitotic events occurred in the anterior of pDCS-treated transplanted planarians, implying asymmetrical effects likely due to pDCS polarity (Fig. 2K). In the sham control group, we did not observe asymmetries in dividing cells within the scarce population of mitotic cells outside transplanted tissues (Fig. 2I-K). We also observed variability in the effects of pDCS over mitotic cells. Some animals showed a low/no response (26%) while the large majority displayed moderate effects (67%) by noticeable mitotic signal outside of the transplant (Figure S1A).

To address the possibility of pDCS induced mitotic asymmetry within irradiated tissues, similar magnitude DCS were applied to 4dpt planarian with opposite polarity (i.e., reversed polarity, implying the positive pole located in the tail and the negative pole in pre-pharyngeal tissue in front of the transplant). These experiments led to inconsistent results, thus we decided to continue with the characterization of pDCS based on polarity with positive pole to the anterior and negative implanted in the posterior region of the animal. We also found pDCS displayed similar effects when the tissue graft was placed in the posterior region (Fig. S1B, C). However, tissue transplants in the anterior region were more reliable and convenient to characterize the pDCS effects.

### pDCS-induced transcription of stem cell markers originates in lethally irradiated tissues

Exposure to a lethal dose of ionizing radiation permanently eliminates neoblasts in less than 24 hours (Peiris et al., 2016a). Distinctively, pDCS leads to the transcription of neoblast markers and the presence of mitotic cells in the host tissue several days after the exposure to lethal ionizing radiation. This finding prompted us to investigate the potential source of neoblast-related cells.

First, since the exogenous application of electric fields is widely known to guide movement of cells through electrotaxis (McCaig et al., 2005), we addressed dynamics of cell migration from the transplanted tissue as the potential mediator of pDCS effects. Previous work determined, neoblasts migrate from transplanted tissue to gradually repopulate the lethally irradiated host at a rate of ∼3-5µm/hr, which is about 72-120 µm/day (Abnave et al., 2017; Eisenhoffer et al., 2008b; Guedelhoefer and Sanchez Alvarado, 2012a; Guedelhoefer and Sanchez Alvarado, 2012b; Newmark and Sánchez Alvarado, 2000; Reddien et al., 2005; Salo and Baguna, 1985). Because neoblasts are the only dividing cells in planarian, the spatiotemporal path of neoblast-related gene expression and mitotic cells (i.e. mitotic wave) are commonly used to infer migration rates between two points (Abnave et al., 2017; Guedelhoefer and Sanchez Alvarado, 2012a; Guedelhoefer and Sanchez Alvarado, 2012b; Newmark and Sánchez Alvarado, 2000; Salo and Baguna, 1985).

Our results show that following 6hrs of pDCS, mitotic cells are found in the tail of the irradiated host, which is ∼5 mm away from the transplanted tissue (Fig. 2I, J). If the transplanted tissue was the source of dividing cells, neoblasts must displace at about 833µm/hr (i.e. ∼200 times faster) to arrive at the tip of the tail more than 700 hours earlier than what has previously been reported (Guedelhoefer and Sanchez Alvarado, 2012a; Salo and Baguna, 1985). Furthermore, we found that shorter pDCS leads to a robust *piwi-1* expression throughout the animal (see results below Fig. 5A with 15 mins pDCS). Were this to be the result of cellular migration from the engrafted tissue, neoblasts must migrate at rates exceeding 20,000µm/hr, or 2cm/hr; which is not only four orders of magnitude faster than previously established but also unlikely due to physical tissue-derived obstacles in their path.

Second, recent findings demonstrate cellular migration in planarians depends on the expression of epithelial-mesenchymal transcription factors *Snail-1*, *Snail-2*, *zeb-1,* and the *ß1-integrin* gene along with components of matrix metalloproteinase (Abnave et al., 2017; Bonar and Petersen, 2017; Isolani et al., 2013; Seebeck et al., 2017). We compared in sham and pDCS treated animals the expression of markers for cellular migration in two segments, the trunk fragment that included transplanted tissue and the tail region at one hour of treatment (Fig. 3A). The results show that in the trunk the expression increased for both *Snail-1*, *Snail-*2 while there was no change for *zeb-1* and the *ß1-integrin*. In the tail region, we found no changes in the expression except for a slight increase in the *ß1-integrin* gene (Fig. 3B, C). Next, we used BrdU to trace migratory cells, but our attempts were unsuccessful due to inconsistent tissue engraftment likely associated with friability of tissue fragments obtained from donors treated with BrdU.

**Fig. 3.**
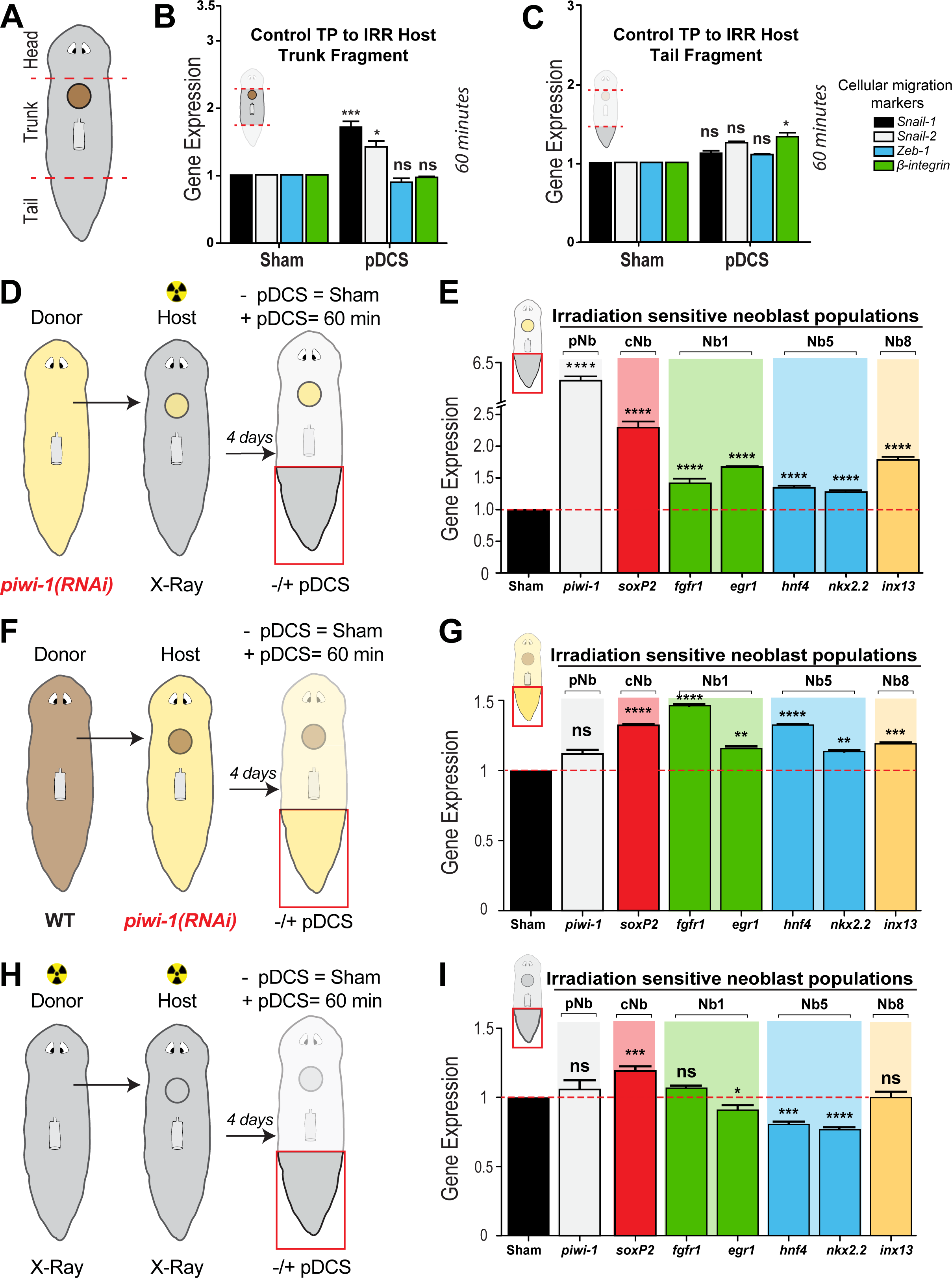
Lethally irradiated host tissue is the main source of pDCS-induced neoblast transcription. (A) Schematic representation depicting different regions of planarians subjected to tissue transplants from wild-type animals. (B-C) Gene expression levels of genes associated with cellular migration in planarians (*Snail-1*, *Snail-2*, *Zeb-1*, *ß1-integrin)*. The tissues used for each experiment were trunk (B) and tail (C) from the sham and pDCS-treated for 60 mins and the gene expression was measured with qPCR. (D, F, H) Depicts experimental design transplanting tissue from different donors and hosts to which tail fragments were processed to measure gene-level expression after four days of transplant. (D) *piwi-1(RNAi)* tissue into a lethally irradiated host, (F) transplant WT tissue into a *piwi-1(RNAi)* host, and (H) transplant γ-irr tissue into a γ-irr host. (E, G, I) levels of gene expression from tissues obtained from sham and planarian subjected to 60 mins pDCS. The gene expression levels involved markers for diverse neoblast populations (*piwi-1*, *soxP-2*, *fgfr-1*, *egr-1*, *hnf-4*, *nkx2.2*, and *inx-13.* Pan neoblast (pNb), clonogenic neoblast (cNb), neoblast (Nb). Data represented as mean ± SEM. All gene expression experiments were obtained from three biological replicates consisting of four-pooled samples per replicate. The polarity of the electric field was positive pole to the anterior and negative to the posterior for 60 minutes. Statistical significance: multiple comparison one-way ANOVA: \**P* ≤ 0.05, \*\**P* ≤ 0.01, \*\*\**P* ≤ 0.001, \*\*\*\**P* ≤ 0.0001.

Third, we did not observe the anterior to a posterior progressive pattern of neoblast expressing cells nor dividing cells that is characteristic during migration-mediated neoblast repopulation of the irradiated host. For instance, stimulation with pDCS using shorter times (i.e., 15 mins) showed strong expression of *piwi-1* at distant places from the transplanted tissue (see results below Fig. 5A at 15 mins). These findings are in stark contrast to the gradual progression of gene expression of *piwi-1* over the AP axis that takes about 40 days to reach the tip of the tail in the absence of electrical stimuli (Abnave et al., 2017; Guedelhoefer and Sanchez Alvarado, 2012b).

Fourth, we designed a series of experiments involving tissue transplantations between, wild type (WT), *piwi-1(RNAi),* and lethally irradiated animals to measure the expression of neoblast markers in the tail region of the host (Figs. 3D, F, H). We performed selective elimination of *piwi-1* expression in either the host or donor tissue to verify the source of *piwi-1^+^* cells and other progenitor subtypes. It is important to note that *piwi-1(RNAi)* specifically silences *piwi-1* expression without affecting neoblast number or function (Reddien et al., 2005). Transplanting tissue from *piwi-1(RNAi)* animal into a lethally γ-irradiated host resulted in a six-fold increase in *piwi-1* expression in the tail of the host subjected to pDCS compared to sham-treated animals (Fig. 3D, E). The increase in gene expression was also prominent in other neoblast markers, suggesting a generalized neoblast response (Fig. 3E). Since *piwi-1* expression was originally silenced in the transplanted tissue, the increased expression of *piwi-1* away from the transplant; specifically, in the tail suggests, *piwi-1* expression originates in host tissues. To confirm this, tissue containing neoblasts from WT animals was transplanted into *piwi-1(RNAi)* host and subjected to identical treatment (Fig. 3F). The results show that *piwi-1* expression was equivalent to sham-treated as expected, but there was an important increase in the expression of other neoblast markers in the tail of animals with pDCS (Fig. 3G). These findings confirm the specificity of the RNAi strategy and provide evidence in support that lethally irradiated host tissue is the source of expression for neoblast markers upon pDCS.

However, it remained unclear whether the presence of neoblast in the graft was needed for the pDCS effects. To address this, neoblasts were eliminated from both the donor and host tissue by lethal γ-irradiation (Fig. 3H). Under these conditions, *piwi-1* expression remained similar to sham control (Fig. 3I), while there was a mixed effect in the expression of markers of neoblast (i.e., most of them were either reduced or did not change except for *sox-P-2,* Fig. 3I). We also noted that application of pDCS in a lethally irradiated animal without transplanted tissue did not trigger expression or neoblast markers or cell division. These results suggest that the presence of neoblasts in transplanted tissue is necessary for pDCS-mediated expression of neoblast markers. Together, our findings indicate that pDCS elicits transcription of SC markers in lethally irradiated tissues, effects that emerge from host tissues but require the presence of grafted neoblasts. Nevertheless, based on the spatio-temporal expression pattern of genes required for migration, the rapid presence of *piwi-1* expressing cells, and cellular division upon short application of electric stimulation, we propose that pDCS-induced effects on neoblast transcription and cell division are not due to electrotactic cellular migration from the transplanted tissue but rather through the activation in the expression of neoblast progenitors and subsequent cell division originating in lethally irradiated host tissue.

### pDCS enhances the DNA damage repair response in tissues exposed to a lethal dose of ionizing radiation

Exposure to a high dose of ionizing radiation induces DNA damage and subsequent cell death (Barghouth et al., 2019; Peiris et al., 2016a; Peiris et al., 2016b; Pellettieri et al., 2010). Nonetheless, 60 minutes of pDCS activates gene transcription, a process known to require DNA integrity. Therefore, we assessed DNA integrity and repair mechanisms on dissociated cells obtained from the lethally irradiated tail region of both the sham-treated and pDCS group. Ionizing radiation increases DNA double-strand breaks (DSBs) that, in planarians, are mainly repaired through homologous recombination (HR) (Barghouth et al., 2019; Peiris et al., 2016b). Immunostainings using markers of DNA damage and repair response were used after 60 mins of treatment in sham and pDCS groups. The results revealed pDCS led to a noticeable increase in phosphorylation of the histone H2AX (γ-H2Ax, Fig. 4A), which is often observed in the early response to DSBs (Bonner et al., 2008; Marti et al., 2006). Likewise, pDCS increased RAD51protein nuclear localization by 20% (Fig. 4B).

**Fig. 4.**
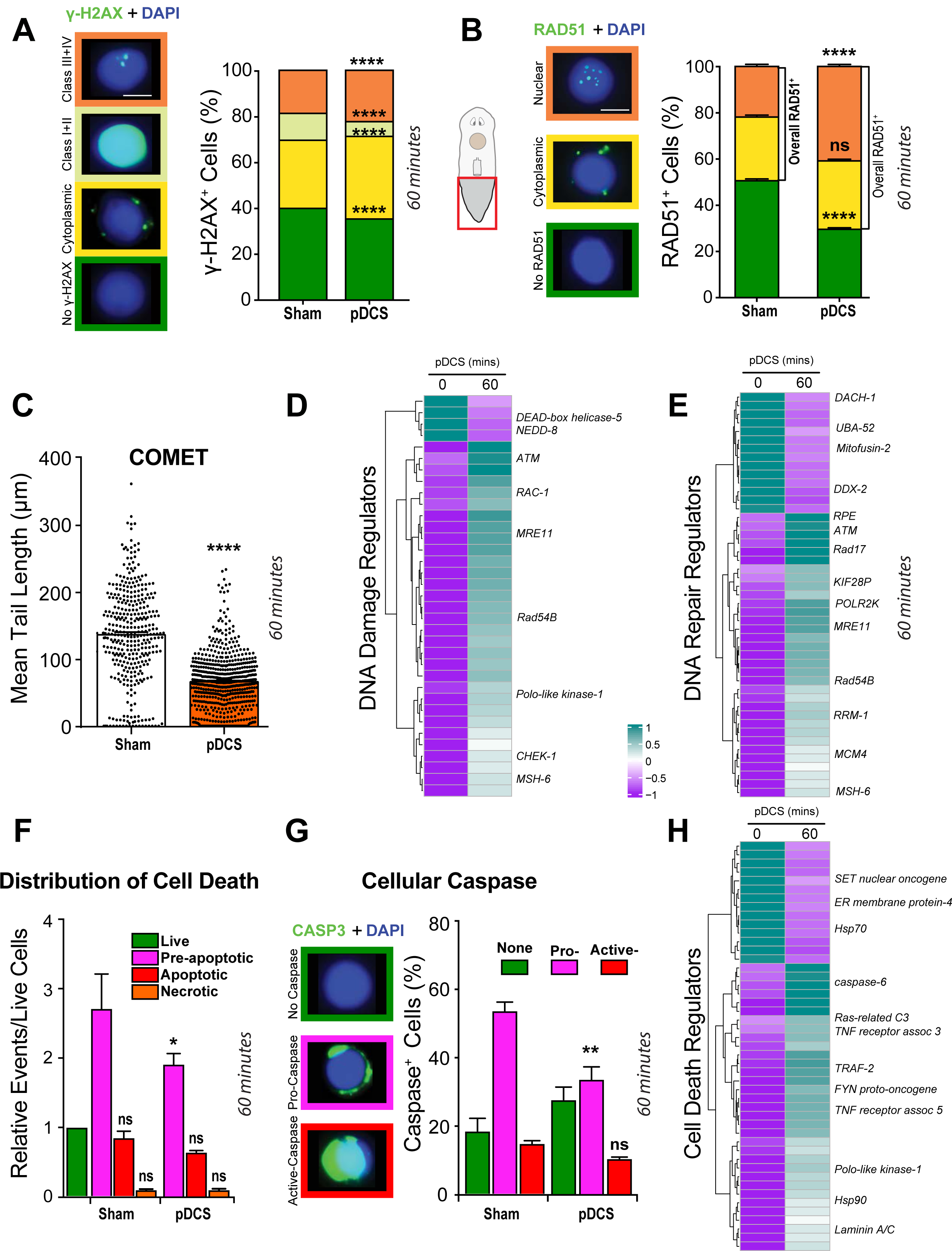
pDCS enhances DNA damage response, reestablish DNA integrity and decreases cell death within γ-irradiated tissues. (A, B) Dissociated cells isolated from tail fragments were immunostained with anti-γH2AX and anti-RAD51 to visualize nuclear (DAPI) vs. cytoplasmic localization in sham and animals subjected to 60min pDCS. Nuclear γH2AX includes four classes of the nuclear signal as displayed in the left of (A) and previously described (Barghouth et al., 2018; Thiruvalluvan et al., 2018). The RAD51 signal was classified based on their localization with respect to DAPI as shown in in the left side of (B). Single-cell extract staining, experiments consisted of five pooled tail fragments and three biological replicates. (C) DNA integrity was measured with the COMET assay (mean tail length) using cells isolated from the host tail fragment in sham and worms subjected to 60min pDCS (n=12 each). (D,E,H) RNA-seq data obtained from tail fragments of sham and 60min pDCS tissue explants (data was collected by pooling tails from four independently treated pDCS or sham planaria across three independent experimental trials). Gene expression heatmaps display differentially expressed transcripts (FDR<0.05) as averaged log2CPM Z-scores for putative DNA damage regulators (D), DNA repair regulators (E), and cell death regulators (H). (F) FACS analysis showing the distribution of live, pre-apoptotic, apoptotic, and necrotic cells using Annexin V in sham and 60min pDCS. The data includes four pooled tail fragments and three biological replicates. (G) Single-cell immunostaining using anti-caspase-3^+^ to denote three possible staining (active, pro-caspase and no caspase images on the left side of (H)). Caspase immunostaining was obtained from five pooled tail fragments and three biological replicates. Data represented as mean ± SEM obtained from experiments independently repeated at least three times. The polarity of the electric field was positive pole to the anterior and negative to the posterior for 60 minutes. Statistical significance: (A-C) Students *t*-test; (F,G) multiple comparison one-way ANOVA: \**P* ≤ 0.05, \*\**P* ≤ 0.01, \*\*\*\**P* ≤ 0.0001.

**Fig. 5.**
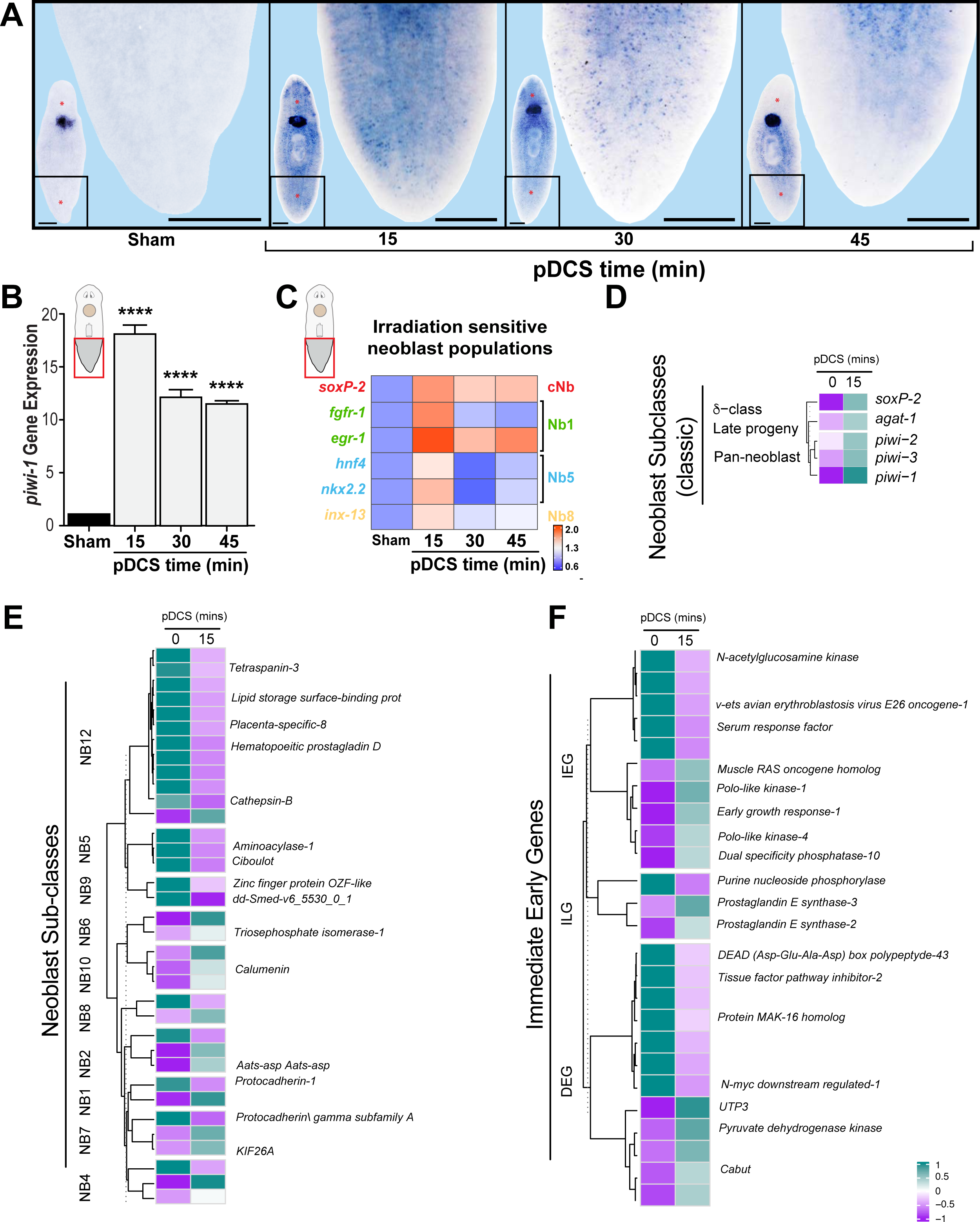
pDCS induces a rapid transcriptional response. (A, B) *piwi-1* gene expression with (A) whole-mount *in situ* hybridization and (B) qPCR at 15, 30, and 45 min of pDCS compared to the sham group. (A) lower-left corner shows the body image of lethally irradiated (60 Gy) animals four days post-transplant tissue from wildtype. The inset denotes the amplified tail section (sham = 10/10, 15min pDCS = 8/10, 30min pDCS = 9/13, 45min pDCS = 10/14). qPCR data represented as mean ± SEM obtained from triplicates consisting of five pooled samples per experiment and repeated two times. (C) Gene expression heatmap of neoblast markers from tail tissues explants, measured by qPCR. Shown are log2FC with column scaled Z-score (data is from six pooled tail fragments/replicates and three biological replicates). (D-F) RNA-seq data collected from tail tissue explants of irradiated and control animals (data was collected by pooling tails from four independently treated pDCS or sham planaria across three independent experimental trials) exposed to 60 min of pDCS. Gene expression heatmaps display differentially expressed transcripts (FDR<0.05) as averaged log_2_CPM Z-scores. (D) differentially expressed classic neoblast subclass populations based on reference (van Wolfswinkel et al., 2014b; Wagner et al., 2012). (E) differentially expressed neoblast subpopulations and their respective lineages based on reference (Zeng et al., 2018b). differentially expressed immediate-early gene putative homologs. The polarity of the electric field is denoted with a positive pole to the anterior and negative to the posterior for 60 minutes. Statistical significance: multiple comparison one-way ANOVA: \*\*\*\**P* ≤ 0.0001. Scale bars, 500µm.

Nuclear translocation is essential for the function of RAD51 during DSB repair (Haaf et al., 1999; Peiris et al., 2016b). We further determined, through comet assay, that pDCS-mediated activation of the DDR was accompanied by a noticeable reduction in DSBs caused by γ-irradiation (Fig. 4C). The results were expanded by performing transcriptomic analysis with a focus on genes associated with DNA damage and repair. RNA was extracted from the tail fragments from sham and pDCS animals after 60 mins of treatment and we used BLAST domains as annotations for the transcriptome dd_Smed_v6 (Grohme et al., 2018). When examining orthologs of DNA damage and repair pathways in *H. sapiens* the analysis confirmed genes associated with these pathways were differentially expressed with a majority upregulated (e.g., *ATM*, *RAD17*, *MRE11*, *Rad54B*) (Fig. 4D, E, Supplemental file 1). A gene enrichment analysis was used to identify the strongly activated pathways after 60 minutes of applied pDCS (Supplemental file 2). The biological processes for DNA damage checkpoints and DNA integrity checkpoints (GO:0000077 and GO:0031570) were considered significantly enriched (KS= 0.0424 and significance 33%, respectively). The biological processes for cellular respiration (GO:0045333), generation of precursor metabolites (GO:0006091), and molecular transport (GO:0008272, GO:0015698, GO:0015698, GO:0072348) were the most highly enriched biological processes. Cellular component genes involved with the nucleus were considered the most enriched cellular components at this time point (Supplemental file 2). This is supported by the enriched molecular functions for nucleotide-binding (GO:0000166), RNA binding (GO:0003723), translation factor activity, RNA binding (GO:0008135), and additional functions involved in nucleic acid activity (Supplemental file 2). The most enriched functions upregulated by pDCS compared to the sham control were catalytic activity (GO:0003824) which showed that 21% of genes annotated with this term were significantly differentially expressed (Supplemental file 2). These data suggest nucleic acid activity is highly enriched upon an hour-long application of pDCS.

In addition to the upregulated DNA repair and DNA damage transcripts, there was a strong upregulation of transcripts associated with replication (Fig. S2, Supplemental file 1). Approximately 60% of significantly differentially expressed genes related to replication were upregulated and further supports the notion that nucleic acid activity is enriched upon application of hour-long pDCS treatment. pDCS effects were also accompanied by improvements in cell viability as determined by flow cytometry with annexin V and immunostaining using Caspase-3 antibody (Peiris et al., 2016a; Thiruvalluvan et al., 2018). We did observe reduced levels of pre-apoptotic cells compared to the sham-treated group (Fig. 4F). Consistently, we also noticed a reduction in pro-caspase-3^+^ cells that are commonly associated with pre-apoptotic cells (Fig. 4G). These results were accompanied by differential expression of genes known to regulate apoptosis (Fig. 4H, Supplemental file 1). In summary, these findings demonstrate that 60 minutes of pDCS is capable of activating DNA repair, DNA damage, and replication mechanisms leading to reduced overall DNA damage in lethally γ-irradiated tissues.

### pDCS activates transcription of immediate early genes in lethally irradiated tissues

Previous work demonstrates the capacity for electric stimulation to produce rapid cellular responses, beginning at the transcriptional level (Dragunow and Robertson, 1987; Saha et al., 2011). To determine if pDCS treatment is capable of influencing transcription of neoblast markers on a more rapid time scale, tissue from WT animals was grafted into lethally irradiated hosts and exposed to different lengths of pDCS (i.e., 15, 30, 45 min; Fig. 5A). Strikingly, expression of *piwi-1* and other neoblast markers were not only detected but found maximally enriched during the first 15 min of pDCS (Fig. 5A-C). The expression of these neoblast-specific genes gradually reduced over time (Fig. 5A-C). Transcriptomic analysis using tail fragments from the 15 mins timepoint showed a strong upregulation in the expression of *Smed-SoxP-2, Smed-Agat-1, piwi-1-3* that are markers of the sigma and pan-neoblast populations according to the classic classification (Fig. 5D, B&H FDR <0.05, Supplemental file 3) (Eisenhoffer et al., 2008a; Kim et al., 2019; Palakodeti et al., 2008a; van Wolfswinkel et al., 2014a). Moreover, we also detected significant differential expression of 35 out of 189 markers (Fig. 5E) associated with various neoblast subclasses (Supplemental file 3) (Zeng et al., 2018a). In general, there was a strong upregulation in the expression of markers associated with cNeoblast (Nb2) and progenitors of the pharynx, (Nb7), epidermal (Nb1), and muscle (Nb4, Nb6) after 15 mins of pDCS (Fig. 5E, B&H FDR= 0.05, Supplemental file 3).

This rapid upregulated transcription pattern was also observed in markers of DNA damage response, DNA repair, and DNA replication (Fig. S3A-C). A gene Set Enrichment Analysis (GSEA) with the topGO Komogorov-Smirnoff test of these data shows the most enriched biological processes occurring after 15 minutes of pDCS are related to metabolic and transport processes (Supplemental file 2). This is also consistent with 40% enrichment of genes related to the mitochondrial outer membrane (GO:0005741). The most enriched molecular function at this early timepoint were genes related to the catalytic activity (GO:0003824) where 21.7% (413) genes were found significantly enriched. RNA binding activity (GO:0003723) genes showed enrichment with 19.6% of genes being significant (Supplemental file 2). Overall, the enrichment analysis suggests DNA damage, repair, and cell cycle processes are activated early upon application of pDCS. Plotting statistically significant genes orthologous to all genes involved in DNA damage, DNA repair, and replication pathways was consistent with the identification of upregulation in critical DNA repair genes. For example, Rad54 and Rad51 are upregulated with a LogFC increase greater than 1.2 (Fig. S4B, B&H FDR= 0.05). Checkpoint Kinase 1 (CHK1) is a critical mediator of DNA damage response and cell cycle activation and upregulated by 1.7-fold with early application of pDCS (B&H FDR= 0.01, moderated t= 5.98). DNA polymerases, helicases, and topoisomerases are upregulated two-fold. Cyclin-dependent kinase 1 (CDK1) exhibits the strongest increase of expression by nearly four-fold and further demonstrates the strong activation of cell cycle pathways (B&H FDR= 0.0046, moderated t= 4.72). In comparison, the expression of CHK1 and CDK1 at the 60 mins timepoint is slightly dampened which supports the high enrichment of cell cycle regulators in the GSEA after 15 minutes pDCS (Supplemental file 2). The results indicate pDCS elicits a rapid transcriptional response geared toward markers of neoblast, DNA damage, and repair within the hosts irradiated tissues.

These remarkably rapid changes in gene expression after pDCS are temporally consistent with activation of immediate-early gene (IEG) transcription, generally defined as genes expressed in the absence of *de novo* protein synthesis (Bahrami and Drablos, 2016; Greer and Greenberg, 2008; Herschman, 1991; Morgan and Curran, 1991; Saha and Dudek, 2013). Furthermore, the mRNA of IEGs are detectable within minutes of exposure to a wide range of stimuli such as stress, mitogens, immune response, neuronal signals, and electric stimulation (Bahrami and Drablos, 2016; Cohen and Greenberg, 2008; Greer and Greenberg, 2008; Saha and Dudek, 2013). pDCS time-course qPCR results show a rapid and transient expression of a well-characterized member of the IEG family, *early growth response gene-1* (*egr-1,* Fig. 5C), which is a well-characterized member of the IEG family (Bahrami and Drablos, 2016; Greer and Greenberg, 2008) and an established neoblast marker required for regeneration and stem cell function in planarians (Lei et al., 2016; Sandmann et al., 2011; Tu et al., 2015a; Wagner et al., 2012; Zeng et al., 2018a).

IEGs are classified based on their induction profile and separated into rapid, delayed, or slow expression response groups following a stimulus (Bahrami and Drablos, 2016; Saha and Dudek, 2013). Thus, we compiled a list of 138 known IEGs previously reported in other model organisms (Figure S4) (Cullingford et al., 2008; Tullai et al., 2007; Uhlitz et al., 2017) and matched the respective expression of their putative planarian orthologs following pDCS. Genes that were considered significantly differentially expressed at the 15mins timepoint were listed following *p*-value and FDR cut-offs of <0.05; this yielded a total of 1778 up-regulated genes. Out of 1778 genes we found 25 planarian orthologs to published IEGs in the 15min pDCS treated planarian (Fig. 5F, S4). This analysis confirmed the upregulated expression of *egr-1* and revealed that the increased expression of IEGs persists through the hour-long application of pDCS (Log2FC= 1.32, B&H FDR= 0.0013, moderated t= 7.42). Additionally, other members of the immediate early genes were considerably upregulated after 15 mins pDCS treatment including the RAS oncogene and the dual-specificity phosphatase 10 (DUSP10) that is known to affect components of the mitogen-activated protein kinases (MAPKs), JNK and ERK (Fig. 5F). Within the 25-planarian putative IEGs we found representation for all three subclasses (IEG, DEG, and ILG) and the number of upregulated and downregulated members of those groups were split. It is not entirely clear how pDCS transcriptional regulation of IEGs is translated into cellular actions, but our observations suggest that short treatment with pDCS stimulates the rapid induction of IEGs within lethally irradiated tissues.

### pDCS induces ectopic expression of neoblast markers mediated by Ca^2+^ signaling

Since exposure to a lethal dose of irradiation irreversibly eliminates neoblasts, we assessed the identity of host-cells expressing neoblast markers following 15 minutes of pDCS. To recognize the spatiotemporal distribution of cells at different stages of differentiation, we performed expression analysis using double fluorescent *in situ* hybridization (FISH) in tissue sections (Fig. 6A). Specific genes were chosen to reflect two distinct stages of cellular differentiation, *piwi-1* and *agat-1*, which label early progenitors and late post-mitotic progeny, respectively (Eisenhoffer et al., 2008b). Our results show that 15 min pDCS induces a 56-fold increase in *piwi-1^+^* cells and this upregulation coincides with a simultaneous 21-fold increase in *agat-1^+^* cells (Fig. 6B, C). Intriguingly, 70.1% of *agat-1^+^*cells co-express *piwi-1* while 24.5% of *piwi-1^+^* cells co-express *agat-1* (Fig. 6B). Moreover, sham control planarian exhibited minimal *piwi-1^+^/agat-1^+^* cells, as expected in 4-day post-irradiated tissue (Fig. 6A, C). Previous research reported minimal, if any, the overlap between *piwi-1* and *agat-1* occurs in WT planarian (< 2.0%) (Eisenhoffer et al., 2008b). The ectopic expression of the neoblast marker in post-mitotic cells suggests that the brief application of pDCS disrupts patterns of gene expression across cellular lineages.

**Fig. 6.**
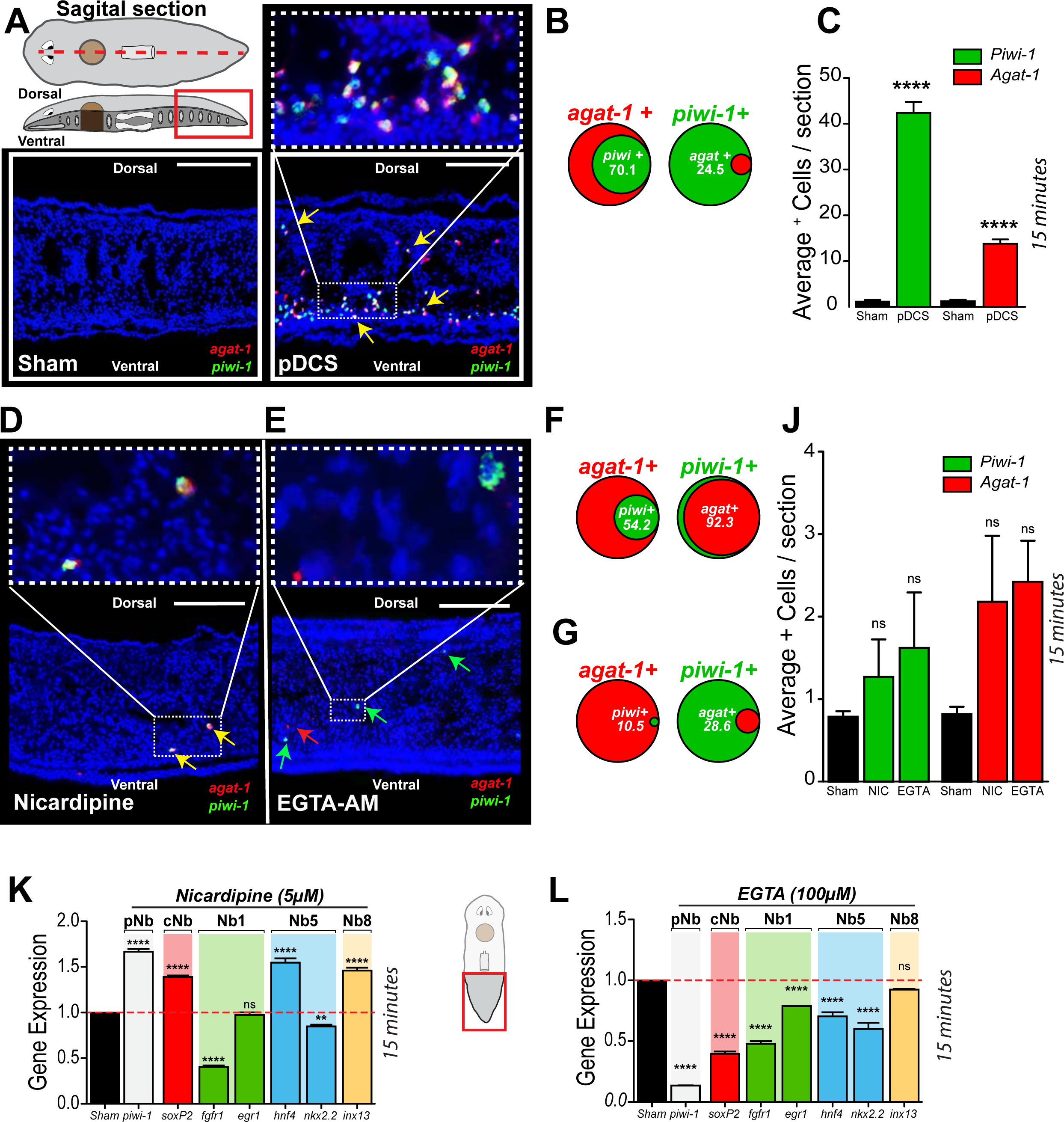
Ca^2+^ signaling mediates pDCS-induced ectopic expression of piwi-1. (A) Double fluorescent *in situ* hybridizations (FISH) were performed on sagittal cross-sections of sham and experimental group (n=six each). (B) Venn diagrams show percentage of *agat-1*^+^ cell population expressing *piwi-1*^+^ (left) or *piwi-1*^+^ cells expressing *agat-1*^+^ (right). (C) The average number of *piwi-1*^+^ and *agat-1*^+^. (D, E) Double FISH performed on sagittal cross-sections following 24hr soak with 5µM nicardipine or 60min incubation with 100µm EGTA [final concentration] (n=six each). (F-G) Venn diagrams show percentage of *agat-1*^+^ cell population expressing *piwi-1*^+^ (left) or *piwi-1*^+^ cells expressing *agat-1*^+^ (right) planarian exposed to 5µM nicardipine or 100µM EGTA. (J) Bar graph displays the average number of *piwi-1*^+^ and *agat-1*^+^ within each inhibition group. (K-L) Gene expression levels of neoblast markers *piwi-1*, *soxP2*, *fgfr1*, *egr1, hnf4, nkx2.2, and inx-13* using qPCR following 24hr 5µM nicardipine soak and 100 µM EGTA (data is from five pooled tail fragments/replicates and three biological replicates). All cases involve isolated tail tissue and pDCS for 15 min in the experimental group. Data represented as mean ± SEM. Pan neoblast (pNb), clonogenic neoblast (cNb), neoblast (Nb). The polarity of the electric field is denoted with a positive pole to the anterior and negative to the posterior for 60 minutes. Statistical significance: (C) Students *t*-test; (F) Kolmogorov-Smirnov test, KS< 0.05 (I, K-L) multiple comparison one-way ANOVA: \*\**P* ≤ 0.01, \*\*\**P* ≤ 0.001, \*\*\*\**P* ≤ 0.0001. Scale bars, 100µm.

Calcium signaling is among the most prominent mediator of excitation-transcription coupling and IEG activation (Greenberg et al., 1986; Greer and Greenberg, 2008; Saha and Dudek, 2013; Saha et al., 2011; Yan et al., 2014). For example, voltage-dependent calcium channels at the plasma membrane can be electrically stimulated to allow the rapid influx of Ca^2+^ to the cytoplasm (Yan et al., 2014). Similarly, calcium signaling mechanisms have been suggested as mediators of acute signaling events in various experimental models, including planarian (Bahrami and Drablos, 2016; Chan et al., 2017; Cohen and Greenberg, 2008; Greenberg et al., 1986; Greer and Greenberg, 2008; Herschman, 1991; Kandel, 2012; Ma and Yan, 2014; Marchant, 2019; Morgan and Curran, 1991; Saha and Dudek, 2013; Saha et al., 2011; West and Greenberg, 2011; Yan et al., 2014). In concert with these reported findings, inhibiting calcium flux through L-type voltage-gated calcium (Cav) channels using a dihydropyridine (DHP), nicardipine, dramatically suppressed the effects of rapid (15 min) pDCS-mediated expression of neoblast markers (Fig. 6D, G, J). The effects of nicardipine inhibition persist even if pDCS was extended to 60 minutes (Fig. S5 A-C). These results were confirmed with nifedipine, another DHP that blocks L-type Cav channels via a different high-affinity binding site (Fig. S5 D-G). Likewise, buffering of intracellular Ca^2+^ with EGTA-AM [ethylene glycol-bis(ß-aminoethyl ether)-N,N,N’,N’-tetraacetoxymethyl ester] also disrupts pDCS-mediated *piwi-1* and *agat-1* expression (Fig. 6H, E, G, K). These results suggest that Ca^2+^ released from intracellular Ca^2+^ stores (i.e., endoplasmic reticulum) mediate pDCS effects.

## DISCUSSION

Our findings underscore the overriding capacity of bioelectric signaling to rapidly affect essential cellular processes such as transcription, cell cycle, and DNA repair. This robust and effective strategy is capable of altering cellular behavior *in situ*, without the need for genetic or pharmacological intervention. The results in planarians are consistent with the overriding effects obtained with DCS in mice models of the Rett syndrome (RTT). The RTT is caused by inactivation of the X-linked gene methyl-CpG-binding protein 2 (*MECP2*) (Amir et al., 1999) and is a complex degenerative dysfunction involving many genes and neuronal groups, in which pharmacotherapy is unlikely to succeed (Baker et al., 2013; Chahrour et al., 2008; Johnson et al., 2017; Sugino et al., 2014). However, application of DCS with electrodes implanted in the brain of RTT mouse model, activate neurogenesis and restore neural circuits and spatial memory, and the behavior of the experimental group is indistinguishable from sham-treated mice (Hao et al., 2015; Lu et al., 2016; Pohodich et al., 2018). Jointly, the results in both vertebrate and invertebrates suggest the overriding effects of DCS (pulsing or steady-state) consistently overcome conditions involving dysfunctional DNA.

We introduce planarians as a simplified platform to carry out comprehensive analysis aimed at dissecting the molecular basis of electric stimulation at the organismal, cellular, and subcellular levels. We observe extensive commonalities between DCS effects in planarians and mammals. For example, the time and strength of the currents used in our DCS are similar to the ones used in humans (e.g. tDCS, muscle, bone repair) (Gerovasili et al., 2009; Kadosh et al., 2010; Moreno-Duarte et al., 2014). Ca^2+^ signaling consistently recurs as a mediator of DCS effects in planarians and mammals. Likewise, the overall changes upon pDCS are transient, thus providing self-contained regulatory mechanisms that can be calibrated to gain desired cellular responses under different circumstances. Uniquely, our findings demonstrate a cost and time-effective alternative to study rapid activation of transcription in tissues exposed to high doses of ionizing radiation. DNA damage is central to cancer, aging, and radiotherapy, but there are limited options to effectively enhance genomic repair. We present evidence demonstrating short exposure to pDCS activates transcription of genes involved in the DDR, which together lead to the reestablishment of DNA integrity in tissues exposed to a high dose of ionizing radiation. Future experiments will be designed to address both the fidelity of pDCS-induced DNA repair and the molecular mechanism mediating this process. One possible candidate may involve small non-coding RNAs (sncRNAs), which recent evidence shows may facilitate the recruitment of repair components in both HR and NHEJ to sites of DSBs (Gao et al., 2014; Qi et al., 2016; Wei et al., 2012).

pDCS triggers the ectopic transcription of stem cell and differentiated tissue markers followed by mitotic activity in tissues damaged by ionizing radiation. The molecular mechanism of this intriguing finding is still unclear, but it is possible that pDCS may affect cell fate regulators associated with the lineage progression of post-mitotic progenitors to increase cellular plasticity. Indeed, our finding showing overlapping expression of *agat-1* and *piwi-1* in post-mitotic cells is consistent with recent findings demonstrating enhanced cellular plasticity by disturbing hippo and *egr-5* signaling pathways (de Sousa et al., 2018; Tu et al., 2015b). However, to our knowledge, there is no precedent in which a short electric stimulus can robustly coordinate genetic and cellular events toward stem cell reconstitution in tissues damaged by ionizing radiation. Future studies will be needed to understand the epistatic relationship between cell fate regulators and the identity of the cells expressing neoblast markers ectopically along with the long-term stability of the *piwi-1^+^* cells.

One possible explanation driving the pDCS-mediated expression of stem cell markers is that distinctive post-mitotic progenitors can sense the electric stimulus and respond. Consistent with this idea, we propose post-mitotic lineages expressing L-type Cav channels [i.e., neural, epidermal, parenchymal, protonephridia, and cathepsin^+^ cells (Fincher et al., 2018; Plass et al., 2018)] are likely the targets of pDCS-enhanced plasticity (Fig. S6). Future experiments will address the individual contribution of post-mitotic lineages expressing L-type Cav channels after pDCS. Another possible scenario may involve the presence of radio-resistant neoblasts or neoblasts with low cycling activity that are sensitive to the electric stimulus. Recent evidence supports the possibility of slow-cycling neoblasts with distinctive regenerative properties (Molinaro et al., 2021). The complete picture of the neoblast heterogeneity and their regulation is an evolving subject that is far from being understood (Fincher et al., 2018; Molinaro and Pearson, 2016; Plass et al., 2018; Reddien, 2018; Rink, 2018; van Wolfswinkel et al., 2014b; Wagner et al., 2011; Zeng et al., 2018b).

We propose a model whereby pDCS may lead to enhanced DNA repair followed by transcription. Our results implicate pDCS effects are mediated by increases in intracellular [Ca^2+^] concentration via L-type Cav channel and/or release from intracellular Ca^2+^ stores (Fig. 7A). The initial effects of pDCS stimulate transcription of IEGs that are tightly regulated by increases in intracellular [Ca^2+^] concentration via L-type Cav channel and/or release from intracellular Ca^2+^ stores (Fig. 7B). The increase in cytosolic [Ca^+2^] may serve to boost DNA repair improving DNA integrity followed by the transcription of IEGs (Fig. 7B). The outcome of these Ca^2+^ mediated events significantly reduce overall levels of DNA damage leading to transcription of stem cell-related genes and cell cycle re-entry in tissues damaged by ionizing radiation. Further experiments are needed to determine the order in which these events take place and to further define the identity of the cells expressing neoblast markers. It is unclear whether radio-resistant cells, slow-cycling neoblasts, or post-mitotic cells with enhanced plasticity are associated with pDCS-induced effects (Fig. 7C).

**Fig. 7.**
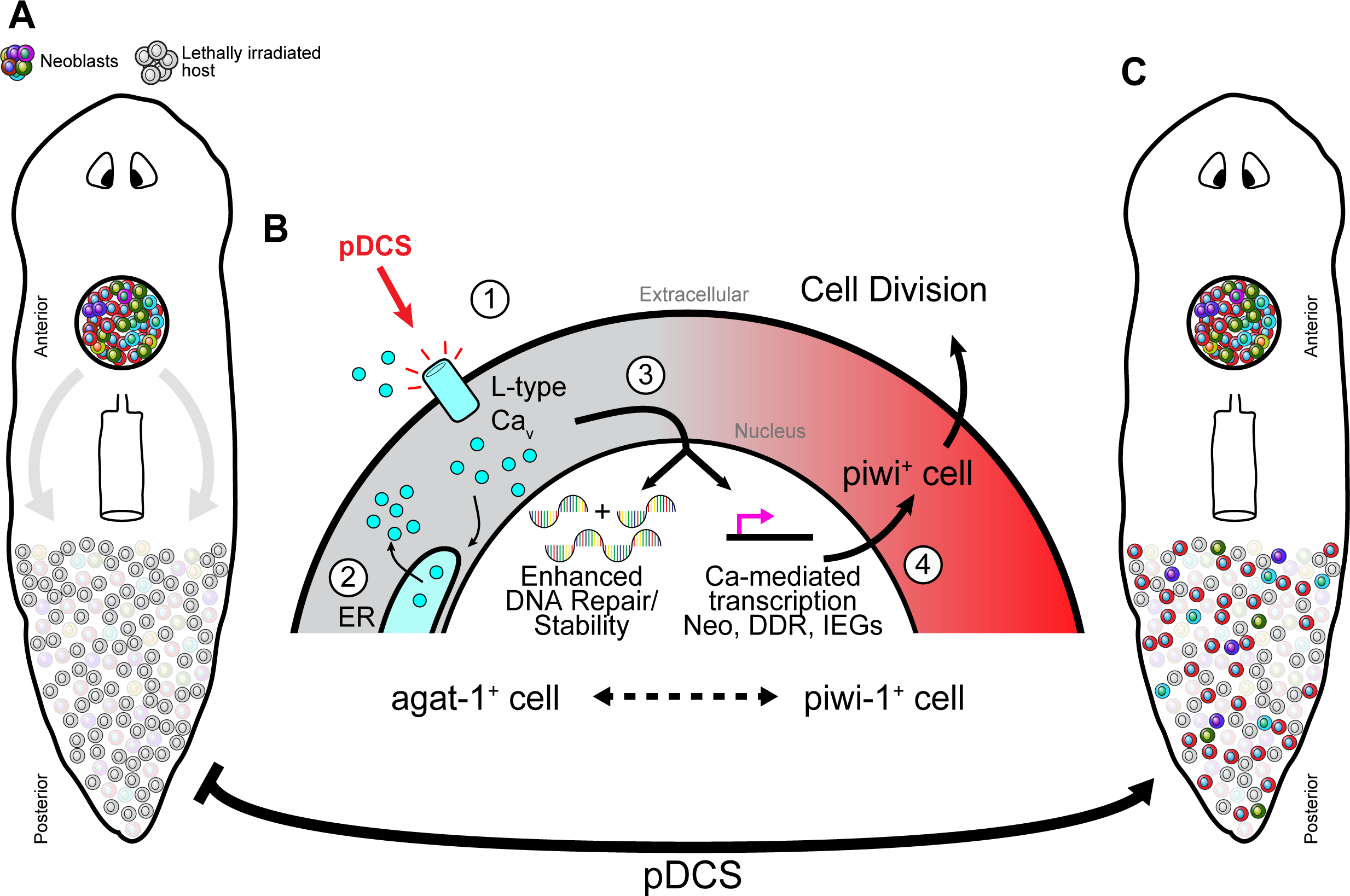
Schematic summary of pDCS-induced effects. (A) pDCS-mediated effects in lethally γ-irradiated host tissue (gray cells) require engrafted neoblasts (colored cells, gray arrows). (B) Proposed cellular effects responsible for observed pDCS-induced transcriptional activation. Transcription sensitive to (1) Ca^2+^ influx through L-type Cav and subsequent (2) Ca^2+^ release from intracellular stores (i.e., endoplasmic reticulum) leading to (3) Ca^2+^-mediated transcription and (4) expression of pDCS-induced genes. The ectopic overlapping expression of *agat-1^+^* populations with *piwi-1* suggests enhanced plasticity of transcriptional programs within the lethally γ-irradiated host tissue. ER=Endoplasmic Reticulum. (C) pDCS activates rapid transcription of genes associated with neoblast populations in lethally γ-irradiated host tissue.

## Material and Methods

### Planarian culture and maintenance

For transplantation experiments, planarians were acclimatized to a final culturing temperature of 13°C. Acclimatization was gradual, beginning with an initial transfer to 16°C for approximately two weeks and then stepped to 13°C permanently until planarian were used for transplantation. Before each change in temperature, planarian cultures, destined for transfer, were fed before temperature changes. Animals transferred to 13°C incubators were cleaned once per day for the first four days of temperature acclimatization. All planarian maintenance was performed as previously described (Oviedo et al., 2008b). After the first four days, planarian maintenance was resumed as previously described (Oviedo et al., 2008b). Planarians were not used for tissue transplantation until they had been cultured for at least two weeks at 13°C. Reduced culturing temperatures were used to decrease the mobility of recovering planarian following tissue transplantation.

### Tissue Transplantations

Planarians were transplanted as previously described (Guedelhoefer and Sanchez Alvarado, 2012a; Guedelhoefer and Sanchez Alvarado, 2012b) with minor changes to tools used for transplanting tissue. We developed a transplantation tool to facilitate consistency in the size of the graft and reduce tissue damage. Briefly, the transplantation tool was made from a 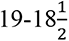 gauge syringe that was bored out to an inner diameter of 750µm using a Dremel drill bit. The outer diameter was polished using 500-1000 grit wet sandpaper until the edges were paper thin and smooth to reduce drag during tissue insertion.

Transplantation schedules varied with respect to experimental condition (i.e. WT, irradiated, or RNAi tissues). Specifically, in experiments using irradiated planarian host or donor tissue, all irradiation was performed 24hours before tissue transplantation. For transplantation experiments using piwi-1 RNAi tissues, transplantation was performed 48hrs post-final injection (5^th^ injection) and subsequent pDCS experiments occurred 4 days post-transplantation. Additionally, in transplantation experiments using *piwi-1* RNAi planarian, all *piwi-1* RNAi donor tissue was derived from *piwi-1* RNAi hosts (i.e. *piwi-1* donor inserts were taken from the anterior region of intact *piwi-1* RNAi who were then used as hosts for wild-type donor tissues).

### Planarian Immobilization

Transplanted or intact planarians were placed in chilled 0.2% w/v chloretone solution for 3-5 minutes (Guedelhoefer and Sanchez Alvarado, 2012a; Guedelhoefer and Sanchez Alvarado, 2012b) in preparation for agar immobilization. After soaking, the planarians were rinsed in chilled planarian water. Motionless planarians were individually placed on large 75mm x 50mm glass slides roughly 1.0 cm apart, atop the ice. All remaining planarian water was removed, and planarian were subsequently covered in 1.0% low melting point agarose (1.0% w/v LMP agar, planarian water) (ThermoFisher 16520050), nearing room temperature, until the planarians were entirely submerged. Planarians were gently positioned to level during the gelation process to achieve maximum body axis symmetry. Once the agar is completely solidified, excess agar was trimmed and encapsulated planarians were placed into the center of chilled 35mm Petri dishes (Corning CLS430588), prefilled halfway with solidified 1.0% agarose (1.0% agar w/v, planarian water) (Sigma A9539). The remaining petri dish volume was filled with 1.0% agarose until the agarose level is flush with the top of the encapsulated planarian. All agar encapsulation processes were performed on ice.

### Administration of current and electric field generation for pDCS experiments

Planarians were immobilized in agarose and subjected to applied currents via current clamped microelectrodes. To deliver the current, a power supply was fashioned using thirty 9V batteries in series sectioned in 45V increments; the power supply is fashioned with a 100K rotary potentiometer to adjust output to the desired voltage.

To deliver the current to the planarians, borosilicate sharp microelectrodes were pulled using the P-97 Flaming/Brown pipette puller (Sutter Instruments P-97). Microelectrodes were filled with 3M KCl and placed vertically in a 3M KCl bath with the pulled tip in solution. To deliver current to the planarians, microelectrodes were bridged with 6-8” long 1/32” I.D. PVC tubing filled with 3M KCl 1.0% agar connected to 3M KCl baths joined to the power supply via Pt electrodes. The current was clamped using an RNX 3/8 1GO 100MΩ resistor (Mouser RNX-3/8-100M) in series with the planarian.

For all pDCS experiments, the power supply output was 70V (7µA delivered to planarian) and immobilized planarian were impaled through their ventral epithelial layer, in the pre-pharyngeal region, proximal to the brain, and the tail tip, with each glass microelectrode. The duration of current administration ranged from minutes to hours depending on the needs of the experiment. The polarity of the electric field in this paper was positive pole in the anterior and negative to the posterior region of the animal.

### Library Preparation and RNA Sequencing

Planarians treated with pDCS and sham transplanted planarian tails were isolated, and RNA was extracted for each sample. pDCS and sham planarian for these experiments were planarians with wild-type donor tissue transplanted into 24 hours irradiated host tissue. Triplicated analysis was performed for each data point that consisted of pooled samples from four planarian tails per replicate. RNA libraries were prepared and sequenced on the Illumina HiSeq 4000 platform at the DNA Technologies Core at the UC Davis Genome Center. Samples were indexed and pooled for multiplexing. All samples were analyzed using a Bioanalyzer for quality control before sequencing.

### RNA Isolation

RNA from tissues was extracted as previously described (Oviedo et al., 2008c). Tail fragments from sham and pDCS animals were placed in Trizol immediately after amputation and RNA was extracted for each sample. Triplicated analysis was performed for each data point that consisted of pooled samples from four planarian tails per replicate.

### Library Preparation and RNA Sequencing

The cDNA sequencing libraries were prepared using an automated system at the UC Davis Technologies Core. All samples were accompanied with quality control (QC) documentation and profiled with Bioanalyzer for QC before sequencing. Poly-A enrichment was used to remove ribosomal RNA contamination and maximize mRNA detected. Samples were indexed and pooled for multiplexing. Using the Illumina HiSeq 4000 platform, paired-end reads were sequenced to a length of 200 bp by the DNA Technologies Core at the UC Davis Genome Center. This generated high-quality RNA-seq data for thorough downstream bioinformatic analysis to detect delicate changes in phenotype. Paired-end sequencing was also used to resolve ambiguous differences with high repeat regions.

### Read Mapping and Gene Expression Analysis

Trimmed fastq files were assessed for quality control and mapped to the recently published complete *Schmidtea mediterranea* genome dd_Smes_g4_1 from the PlanMine database (Grohme et al., 2018). The Bioconductor package Rsubread version 2.4.2 (Liao et al., 2013) was used to map reads to the reference genome using a robust and efficient seed-and-vote algorithm followed by the featureCounts algorithm to assign counts. The raw counts’ data were normalized and filtered for genes with log2 counts-per-million (CPM) greater than 0.5 (Supplemental file 4). The sample variation was assessed for quality. A customized pipeline using the limma package and voom transformation for precision weights was developed (Law et al., 2014; Phipson et al., 2016; Ritchie et al., 2015). Limma version 3.46.0 and edgeR version 3.32.1 were used for the statistical analysis. Test statistics were produced using empirical Bayes moderation, and subsequent heatmaps were made using the ComplexHeatmap Bioconductor package. We separately tested the contrasts of gene expression in the 15-minute and 60-minute conditions against that in the sham control using a Benjamini-Hochberg False Discovery Rate of 5%. All bioinformatic analyses were coded with R version 4.0.3 on macOS Big Sur 10.16 using the x86_64-apple-darwin17.0 platform.

### Gene Set Enrichment Analysis

Gene Set Enrichment Analysis was performed using the Bioconductor topGO package version 2.42.0. (Alexa and Rahnenfuhrer, 2020; Alexa et al., 2006). Gene Ontology annotations for the dd_Smed_v6 transcriptome were mined from the Planmine database and used to map pathway enrichment (Alexa and Rahnenfuhrer, 2020; Grohme et al., 2018). Gene rankings calculated by the Limma-voom pipeline were used for determining the significance and enrichment (Alexa et al., 2006; Law et al., 2014; Liu et al., 2015; Phipson et al., 2016; Ritchie et al., 2015; Smyth and Speed, 2003). Enrichment was computed by ranking gene scores using the conservative Kolmogorov-Smirnov (K-S) test. Pathways were considered enriched with a K-S *p-*value <0.05. A complete RMarkdown-based notebook of code to reproduce the transcriptomic and GSE Analysis is available in Supplementary Online Materials.

### Ca^2+^ inhibition via Nicardipine, Nifedipine, and EGTA-AM

Nicardipine and nifedipine inhibition was performed on transplanted planarian 3 days post-transplantation such that the 24hr incubation period concluded at 4 days post-transplantation. Each dihydropyridine was dissolved in 100% DMSO at a concentration of 10mM and then diluted in 50mL of planarian water to a final working concentration of 5µM, as described previously (Nogi et al., 2009). EGTA-AM administration as achieved via posterior injections 1hr prior to pDCS delivery. Firstly, planarian volume was estimated using 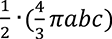 where a, b, and c represent planarian length, width, and height. With estimated planarian volume, EGTA-AM was diluted in Milli-Q water and injected (37nL per injection pulse) such that the final EGTA concentration was 100µM.

### Tissue preparation for cryosections

Planarian are fixed using NAC-formaldehyde based fixation as described previously (King and Newmark, 2013). Fixed animals were prepared for cryosectioning as previously described (Reddien et al., 2005). Briefly, planarians were immersed in increasing concentrations of sucrose diluted in 1XPBS. 1XPBS was replaced with 15% sucrose solution for 1 hour and then 30% sucrose solution overnight at 4°C. Planarians were then placed in tissue embedding molds; 30% sucrose was removed and replaced with optimal cutting temperature (OCT) medium. Planarians were then situated to the desired position/orientation within the OCT and molds were quickly placed in cooling bins containing dry ice immersed in 2-methyl-1-butanol. Once frozen, samples were stored at -80°C until needed for sectioning. The tissue was sectioned in a Leica Cryostat CM1860. Sectioned tissue was either used immediately for FISH/IHC or stored long-term at -80°C until needed.

### Fluorescent *in situ* hybridization (FISH) on sectioned tissue

Cryo-section containing slides were removed from storage at -80°C and given 30min to reach room temperature. Clear scotch tape was placed over the frosted label to ensure label longevity and assist in coverslip placement during hybridization. Custom-made multi-slide chambers were developed and used throughout the procedure. Multi-slide chambers held 12 slides and only required 16ml of the solution to reach maximal coverage. Standard Coplin Jars may be used but require more solution volume per slide contained. After slides reached room temperature, slides were placed in the chamber and rehydrated in 1XPBS for 15min (x2). Following rehydration, FISH was performed as described in (King and Newmark, 2013) with specific changes, as follows: proteinase K incubation was performed for 10min at room temperature at 1µg/mL PK concentration. 1:1 prehybridization/PBStx was omitted. For pre-hybridization and hybridization solution incubation, slides were removed from the chamber and placed in repurposed slide holding containers (converted to hybridization chambers using water-saturated tissue paper placed in each column (x2)) 150µL of the solution was administered to slides, and then glass coverslips were placed atop slides to seal in solution, decreasing evaporation. At the end of the FISH procedure, DAPI (1:1000) was added to slides within the multi-slide chamber for 30min and washed (x2) with 1XPBS. Slides were mounted using the Gelvatol solution.

### Fixation of large planarian for in-situ hybridization

All large planarian (8-12mm) used for whole-mount *in-situ* hybridization (WISH) were fixed using a modified NAC fixation protocol previously described (Guedelhoefer and Sanchez Alvarado, 2012a). Changes for fixation were introduced as follows: planarian first underwent MgCl2 tissue relaxation as described in (Forsthoefel et al., 2014). Briefly, planarians were placed in 0.66M MgCl2 for 45-60s until planarian fully relaxed. The MgCl2 solution was replaced with 10% N-Acetyl cysteine for 10min at room temperature. Lastly, bleaching was performed with a modified bleaching solution (0.5% formamide, 0.36% H2O2, 0.05% Triton X-100, 1X PBS) and placed under light, overnight.

### Whole-mount in-situ hybridization (WISH)

WISH was performed following a previous protocol (Guedelhoefer and Sanchez Alvarado, 2012a). Minor modifications were made during proteinase K treatment, doubling the concentration to 2mg/mL, treating samples for 10min at room temperature.

### Whole-mount immunohistochemistry (IHC) and analysis

Planarians were fixed for IHC following standard formaldehyde-based fixation with an added formamide based step as described previously (Guedelhoefer and Sanchez Alvarado, 2012a). Cells counted positive for Histone H3 phosphate signal (H3P) were normalized against the planarians surface area (mm^2^). When counting H3P in transplanted planarian, the anterior-posterior axis margin was shifted to the center of the transplant. Total H3P^+^ cells were then counted in relation to the population of H3P^+^ cells within the transplant. Briefly, total H3P^+^ cells were counted in host tissue, specifically excluding cells within the transplanted tissue, to H3P^+^ cells restricted in the anterior or posterior regions within sham and pDCS planarian independently (% cells outside transplant). Cell counting was performed using NIH ImageJ cell counter plugin, and all data analysis was performed in Prism.

### Fixation and IHC on dissociated cells

Planarian tail portions were separated and homogenized following pDCS. Homogenate was suspended in calcium and magnesium-free media (CMF) and placed on ice. Cell density was quantified using hemocytometers and cells were plated at 1million/cm^2^ onto glass coverslips. Cells were given 1hour to adhere to the surface and were then fixed with Carnoy’s solution for 2hours on ice. IHC was performed using the human anti-RAD51 antibody and γH2AX antibody as previously shown (Peiris et al., 2016b; Thiruvalluvan et al., 2018).

### RNAi experiments

RNAi was carried out via dsRNA micro-injection (Oviedo et al., 2008a). *piwi-1* and *nelfB* RNAi consisted of 3 consecutive injections followed by one weekly injection until animals were utilized for experimentation (14days post first injection). Injections were administered to the prepharyngeal regions and, due to the size of the planarian (8-12mm), each planarian was given 6-8 pulses of dsRNA 37nL each. *piwi-1* gene was selected and identified using the SmedGD database (Robb et al., 2015). dsRNA was synthesized in vitro as previously described (Oviedo et al., 2008a).

### Quantitative real-time PCR

Quantitative real-time PCR was performed as described (Peiris et al., 2016b). The ubiquitously expressed gene H.55.12e was used as a reference control. Experiments were conducted in triplicates for each condition. Each qPCR experiment was conducted independently at least two times. All qPCR experiments used RNA extractions from tails of pDCS and sham transplanted planarian unless otherwise specified (i.e. qPCR for migratory related genes used tissue sections across the planarian as described in the text). Extracted RNA was then converted to cDNA using the Verso cDNA synthesis kit (ThermoFisher AB1453A). Gene expression is expressed in fold change in comparison to the given control condition.

### FACS Analysis and Comet assay

FACS and comet analyses were performed as previously described (Peiris et al., 2016a; Peiris et al., 2016b).

### Imaging and data processing

All images were captured using a Nikon AZ-100 multi zoom microscope equipped with NIS Elements AR 3.2 software. The brightness and contrast of digital images captured in NIS Elements software were further adjusted in Photoshop (Adobe). Area calculations and cellular foci quantification were carried out using NIH ImageJ software. Mitotic counts were normalized against the planarians’ surface area using ImageJ.

### Statistical Analysis

Data are expressed as the mean ± standard error of the mean (SEM) or fold change ± SEM. Statistical analysis was performed in Prism 2015 software, Graphpad Inc.

## Data Availability

All raw and processed data files associated with this study have been deposited to the NCBI Sequence Read Archive (SRA) submission number SUB8831617. The bioinformatic and RNA-seq analyses pipeline with metadata files are found on the Github repository at: https://github.com/mlegro/RNA-seq-of-pDCS.

## Acknowledgments

We thank Edelweiss Pfister for lab managing and planarian maintenance, and members of the Oviedo lab for insightful discussions and comments on the manuscript. We are grateful to Ivy Pham for assistance with the planarian recovery experiments upon immobilization and Dr. Richard Nuccitelli for advice and critical reading of the manuscript. The sequencing was carried out by the DNA Technologies, and Expression Analysis Cores at the UC Davis Genome Center, supported by NIH shared instrumentation grant 1S10OD010786-01. We thank Monica Britton and Blythe Durbin-Johnson of the UC Davis Bioinformatics Core facility for advice with transcriptomic analysis. This work was supported by the National Science Foundation (NSF) graduate fellowship award 1744620 to EIM, and the University of California Cancer Research Coordinating Committee (Award# CRR-18-525108), and the National Institutes of Health (NIH) National Institute of General Medical Sciences (NIGMS) award R01GM132753 to N.J.O.

## Author contributions

D.D., M.L., P.G.B., A.L.E., and N.J.O. Conceived, designed, and interpreted experiments. D.D., M.L., P.G.B., S.R., B.Z., E.M., D.A., and N.J.O., performed all experiments, acquired and analyzed data. D.D. and N.J.O. wrote the manuscript. All authors read the manuscript, provided comments, and approved the final version.

## Declaration of interests

The authors declare no competing or financial interest.

## Supplemental Information

**Fig. S1.**
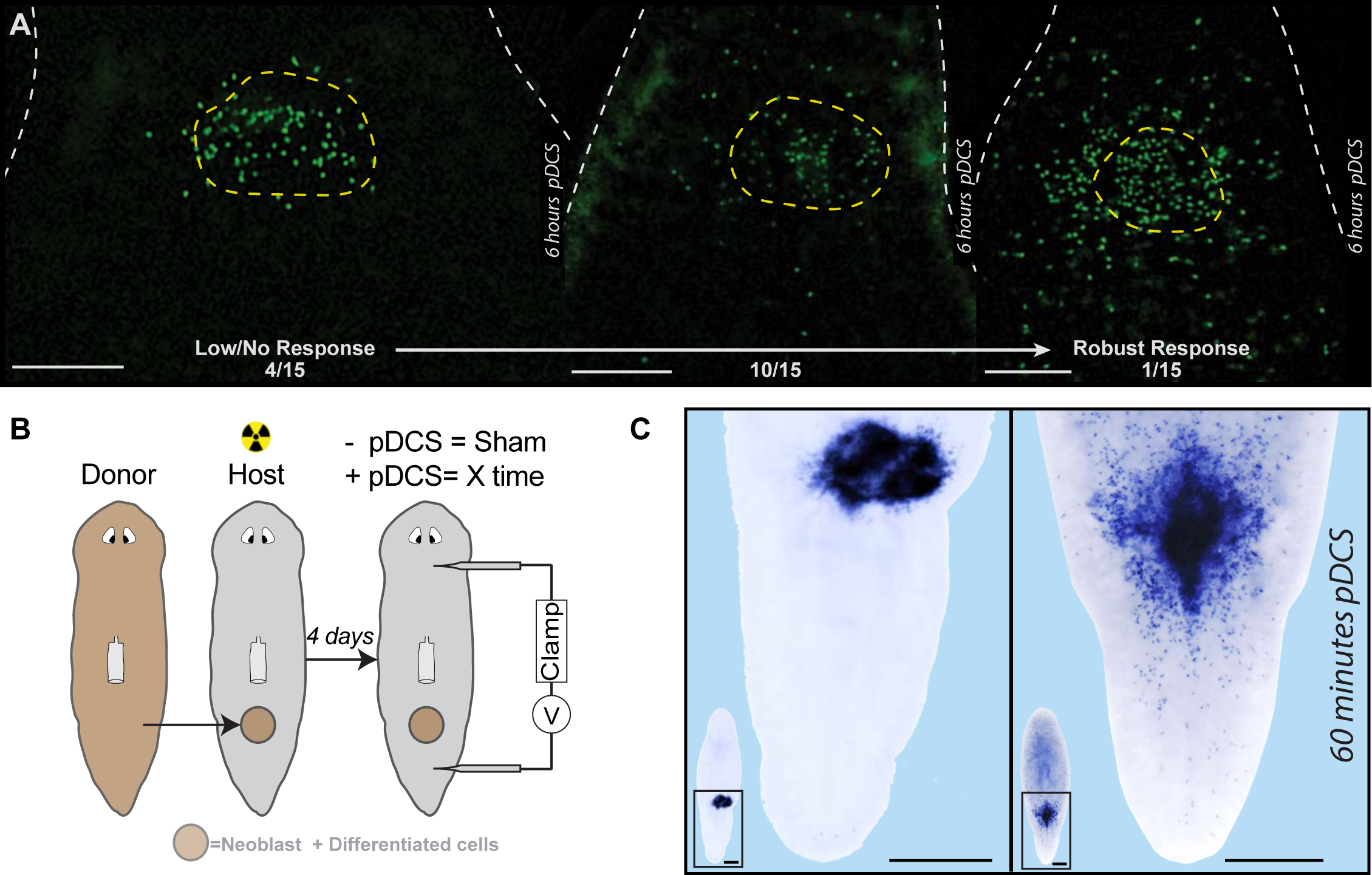
pDCS effects on mitotic activity and transcriptional response in tail transplanted tissues. (A) Immunostaining with anti-H3P antibody (green dots) representing three scenarios where pDCS was able to induce low or no change, moderate and robust number of mitotic cells exiting the transplant after 6hrs pDCS. Number of animals for each condition are included at the bottom of each image. (B) Illustration of current-clamp circuit created by pDCS electrodes and posterior transplanted planarian. (C) WISH addressing *piwi-1* gene expression of 60-minute pDCS. Tissue transplants involved transplant from WT into a γ-irr (60 Gy) host. Scale bars, 500µm.

**Fig. S2.**
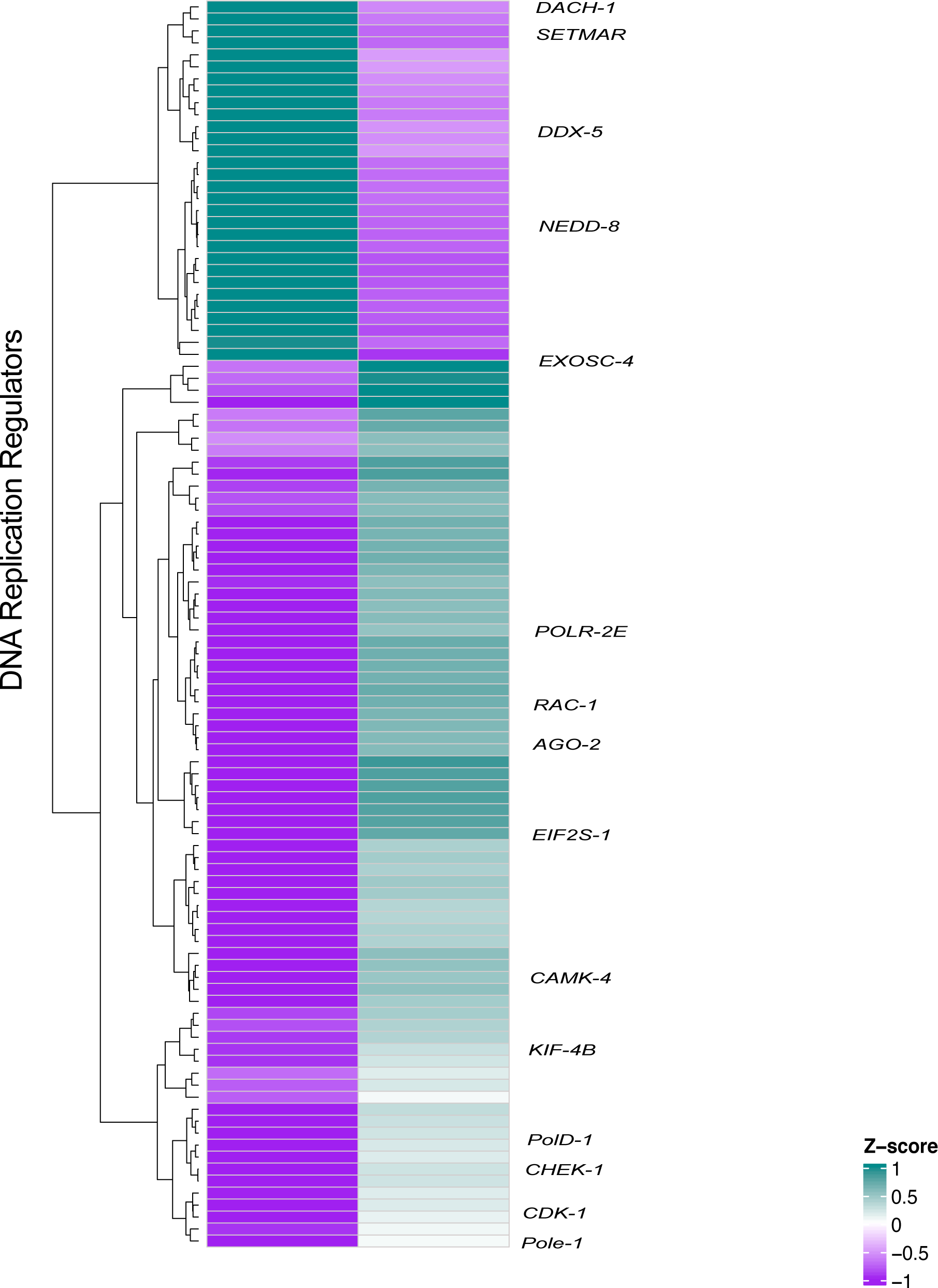
pDCS treatment amplifies DNA replication regulators. Heatmap showing significantly differentially expressed transcripts involved in DNA replication. RNA-seq data collected from tail tissue explants of sham and exposed to 60 min of pDCS (data was collected by pooling tails from four independently treated pDCS or sham planaria across three independent experimental trials). Tissue transplants involved transplant from WT into a γ-irr (60 Gy) host. Gene expression heatmaps display differentially expressed transcripts (FDR<0.05) as averaged log2CPM Z-scores.

**Fig. S3.**
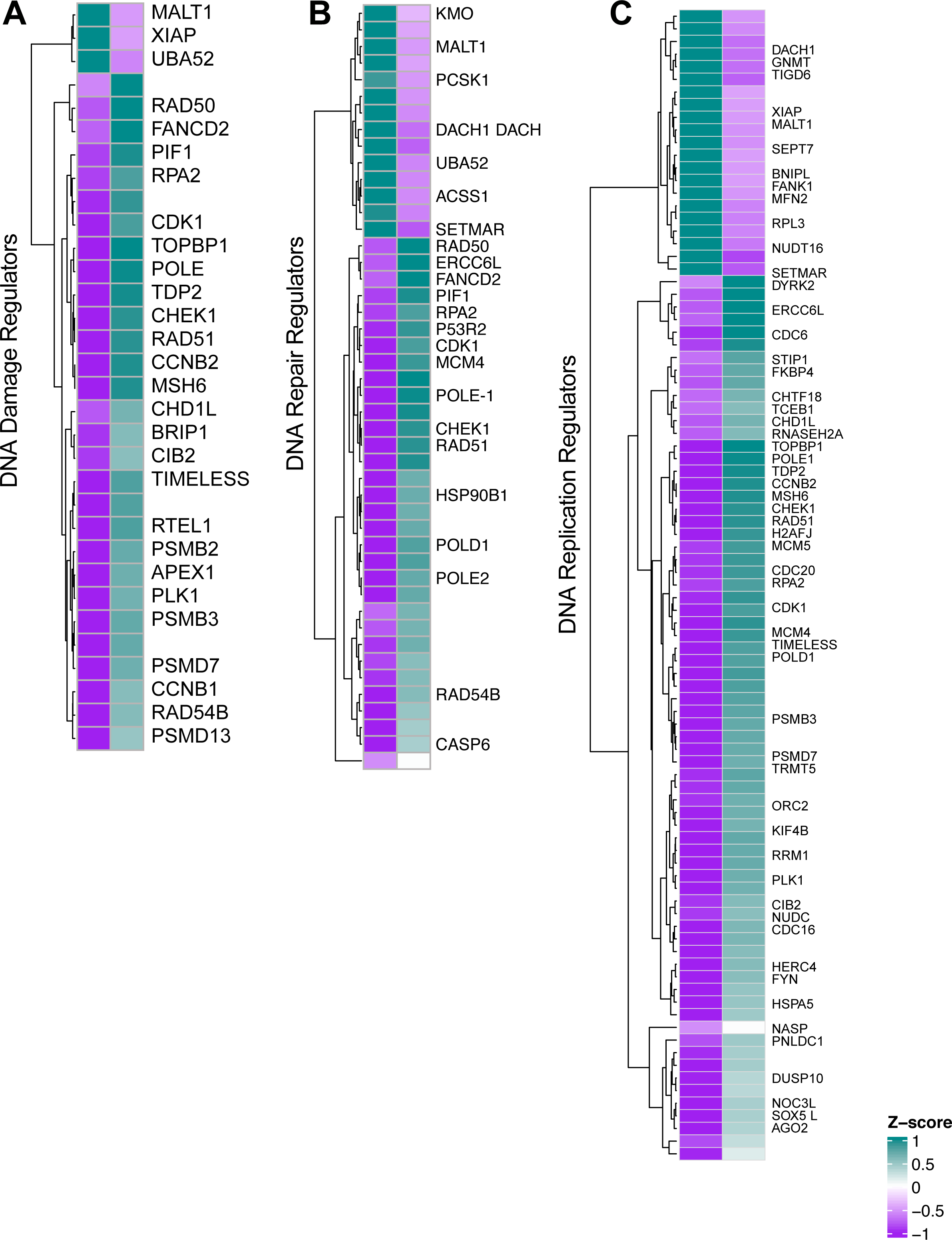
pDCS treatment increases the transcriptional activity of DNA repair, damage, and replication pathways. (A) Heatmaps showing significantly differentially expressed transcripts involved in DNA repair, (B) DNA damage, and, (C) DNA replication. RNA-seq data collected from tail tissue explants of sham and exposed to 60 min of pDCS (data was collected by pooling tails from 4 independently treated pDCS or sham planaria across 3 independent experimental trials). Tissue transplants involved transplant from WT into a γ-irr (60 Gy) host. Gene expression heatmaps display differentially expressed transcripts (FDR<0.05) as averaged log2CPM Z-scores.

**Fig. S4.**
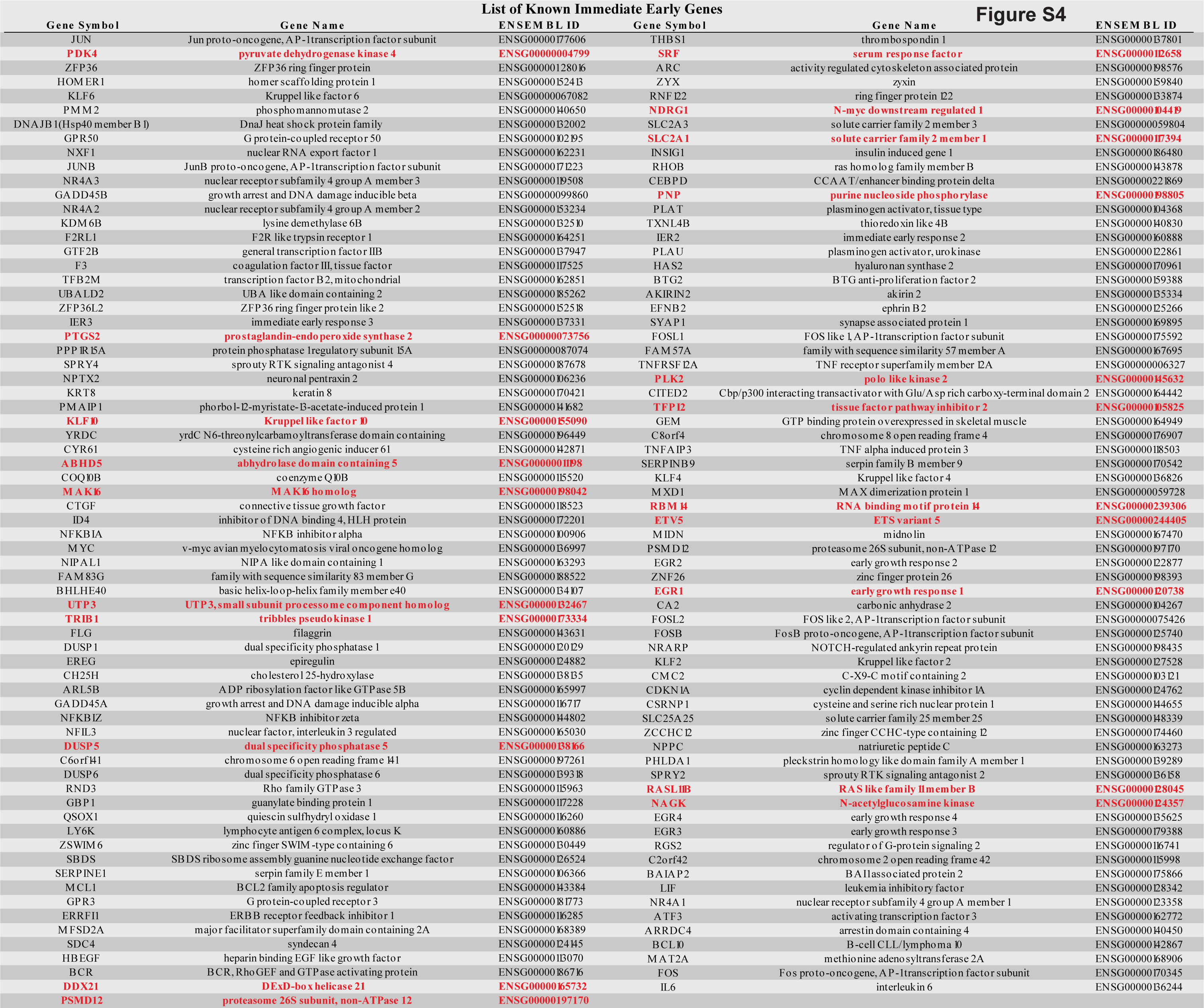
pDCS triggers transcription of immediate early genes. List of previously published and characterized immediate early genes (IEG) from Cullingford et al., 2008; Tullai et al., 2007; Uhlitz et al., 2017. Each IEG is listed with its full name and Ensembl gene ID. Gene names highlighted in red represent planarian differentially expressed putative IEG homologs to vertebrate counterpart.

**Fig. S5.**
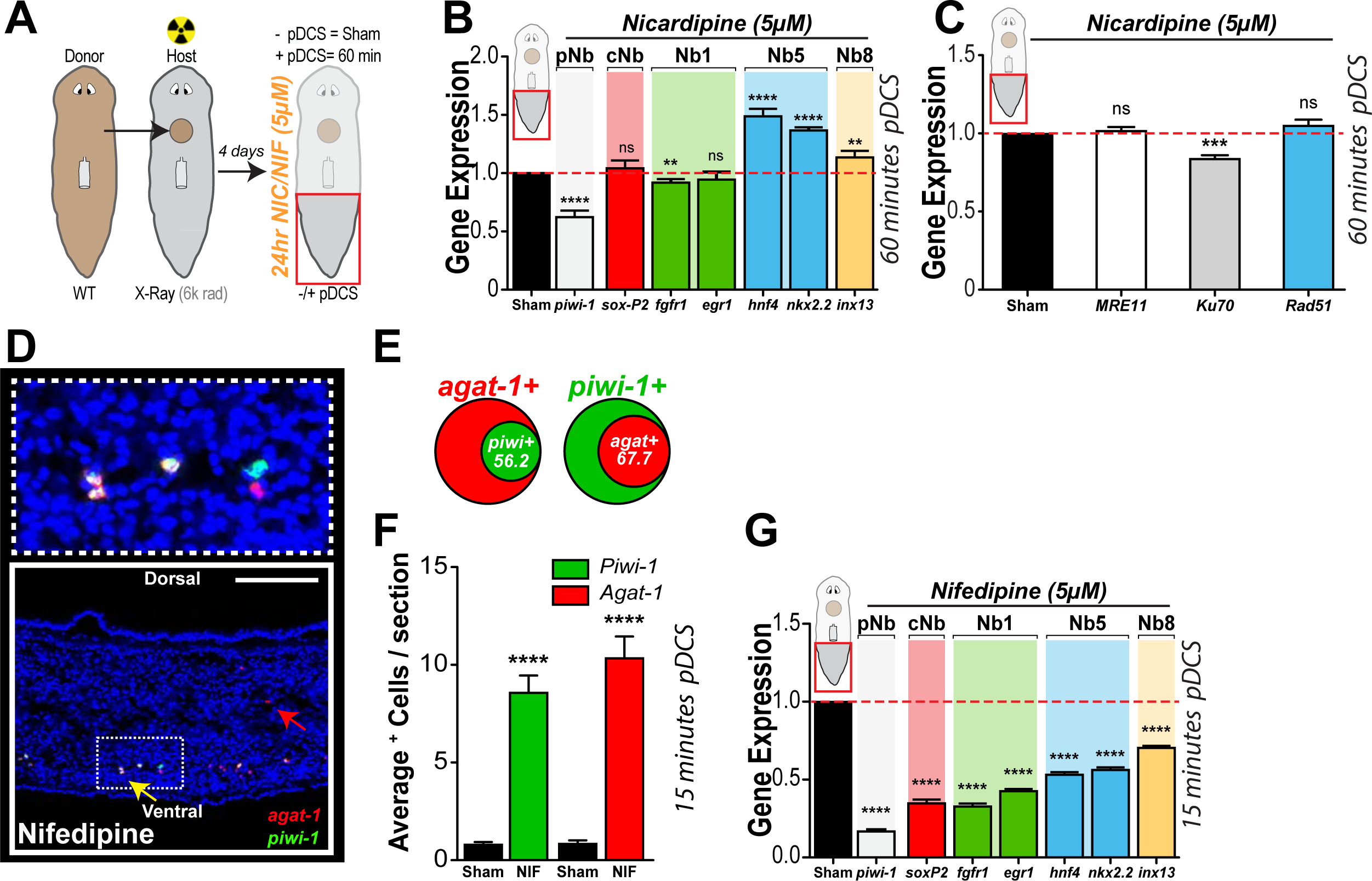
Treatment with dihydropyridines blocks pDCS effects. (A) Schematic showing WT donor/γ-irr host transplanted planarian subjected to 60min pDCS with 24hr 5µM nicardipine soak. Tail tissue was isolated and gene expression for (B) *piwi-1*, *soxP-2*, *fgfr-1*, *egr-1*, *hnf-4*, *nkx2.2*, and *inx-13* as well as (C) DDR genes: *MRE11, Ku70,* and *Rad51* measured by qRT-PCR (n=12, 3 biological replicates). (D) Double FISH was performed on sagittal cross-sections of sham control planarian and planarian exposed to 15min pDCS following 24hr 5µM nifedipine incubation (n=6). (E) Venn diagrams show percentage of *agat-1*^+^ cell population expressing *piwi-1*^+^ (left) or *piwi-1*^+^ cells expressing *agat-1*^+^ (right). (F) The average number of piwi-1^+^ cells and agat-1^+^ cells within tail tissue following 15min pDCS vs. sham planarian per section. (G) Gene expression levels of neoblast markers following 24hr 5µM nifedipine soak measured by qRT-PCR within 15min pDCS compared to the sham group (n=12, three biological replicates). (H) Heatmaps showing significantly differentially expressed transcripts involved in calcium signaling pathways for planarians treated with 15-minutes and 60-minutes pDCS treatment. RNA-seq data collected from tail tissue explants of irradiated and control animals (data was collected by pooling tails from four independently treated pDCS or sham planaria across three independent experimental trials). Gene expression heatmaps display differentially expressed transcripts (FDR<0.05) as averaged log2CPM Z-scores. Data represented as mean ± SEM. Statistical significance: (B, C, G) multiple comparison one-way ANOVA; (F) Students *t*-test: \*\**P* ≤ 0.01, \*\*\**P* ≤ 0.001, \*\*\*\**P* ≤ 0.0001. Scale bars, 100µm.

**Fig. S6.**
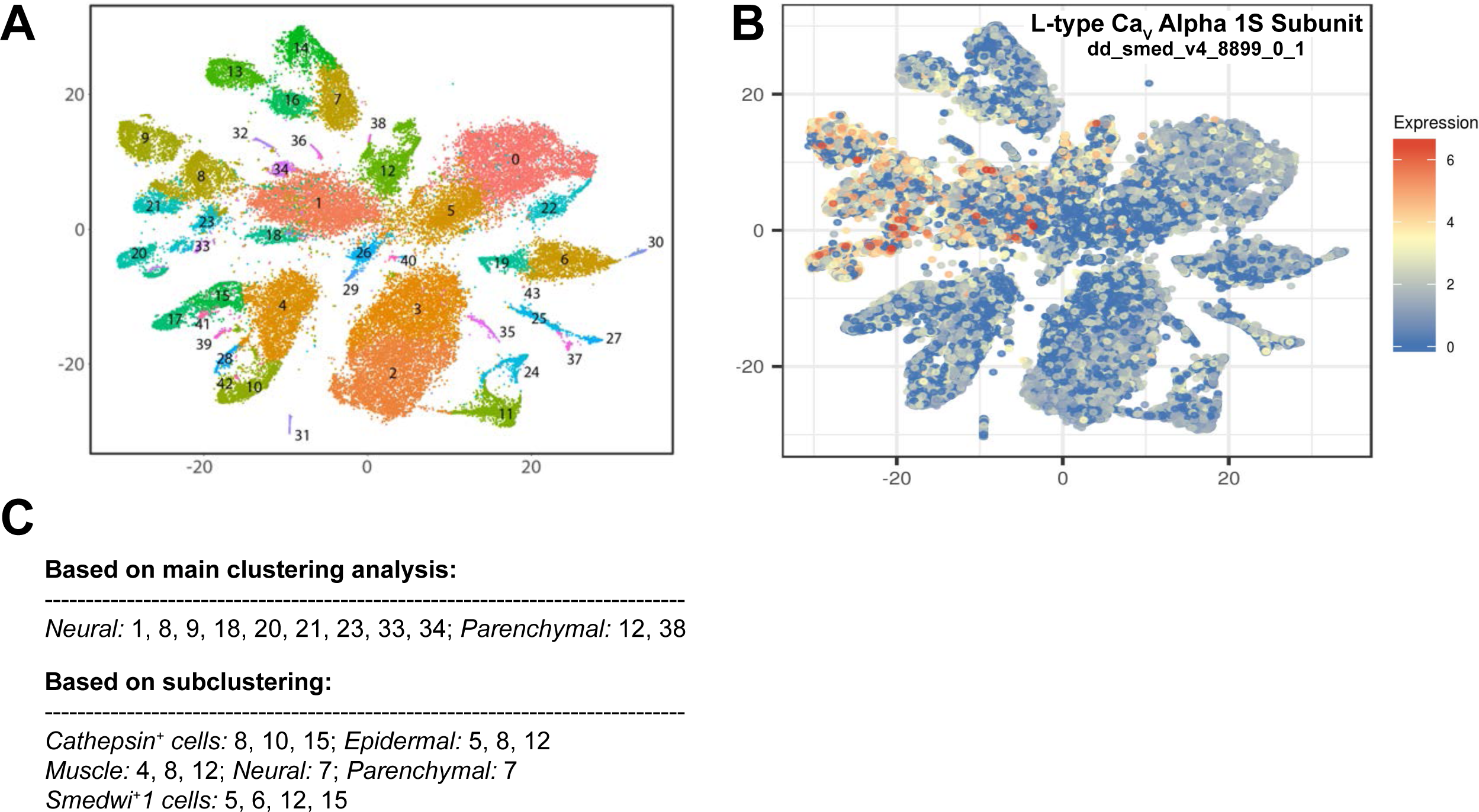
Single-cell transcriptomic analysis displays an extensive spread enrichment of voltage-gated calcium channel alpha1S subunit in planarian tissues. (A) Single-cell distribution of L-type Cav alpha 1S subunit transcript (dd_smed_v4_8899_0_1) taken from *Planarian digiworm* database (Fincher et al., 2018). (B) Graphical key depicting localization for cell populations. (C) Listed cluster enrichment of L-type Cav alpha 1S subunit transcript showing high expression in neural, muscle, parenchymal, epidermal, and cathepsin+ cells.

**Supplemental File 1. Differentially expressed transcripts and associated data for molecular pathway heatmaps.** RNA-seq data showing differentially expressed transcripts for the 15 and 60 minutes timepoints and the associated BLAST data for *Homo sapiens* orthologs. Lists for cell cycle, DNA damage, DNA repair, DNA replication, and cell death molecular pathways are in the associated tabs of the excel workbook. All transcripts are expressed below the 5% level (B&H FDR<0.05) as averaged log2CPM Z-scores.

**Supplemental File 2. Enriched biological processes, cellular components, and molecular functions after applied pDCS.** Gene set enrichment analysis of RNA-seq data collected from the posterior regions of sham and planarians treated with 15 and 60 minutes of pDCS. Tissue transplants involved transplant from WT into a γ-irr (60 Gy) host. Tables plot the most significantly enriched Biological Processes, Cellular Components, and Molecular Functions. RNA-seq data collected from tail tissue explants of irradiated and sham animals (data was collected by pooling tails from four independently treated pDCS or sham planaria across three independent experimental trials). Kolmogorov-Smirnov test, KS< 0.05.

**Supplemental File 3. Differentially expressed transcripts and associated data for NBSC and IEG heatmaps.** RNA-seq data showing differentially expressed transcripts for the 15 and 60 minutes timepoints and the associated BLAST data for transcript orthologs. Lists for NBSC (Reddien, 2018; Rink, 2018; Zeng et al., 2018b; Zhu and Pearson, 2016) and IEGs are in the associated tabs of the excel workbook. All transcripts are expressed below the 5% level (B&H FDR<0.05) as averaged log2CPM Z-scores.

**Supplemental File 4. RNA-seq analysis statistical results for all transcripts.** RNA-seq analysis statistical results the 15 and 60 minutes timepoints. Lists include the toptables of test statistics and CPM Z-score values for all 18,419 transcripts computed in the analysis. This list excludes lowly expressed and filtered transcripts. All transcripts are expressed as averaged log2CPM Z-scores.

## Notes

### Competing Interest Statement

The authors have declared no competing interest.

